# Unsupervised Variant Clustering Identifies Genetic Subtypes of Disease

**DOI:** 10.1101/2025.07.01.662666

**Authors:** Dalia Mizikovsky, Naomi R. Wray, Sonia Shah, Nathan Palpant

## Abstract

Current approaches to identify disease subtypes rely on prior knowledge or external phenotypic references that limit their generalisability across traits. This study presents an unsupervised, phenotype-free framework that analyses the genetic co-occurrence of variants across individuals to identify variant clusters underpinning disease heterogeneity. The approach is developed and validated using simulated combinations of real phenotypes then applied to dissect disease subtypes of type 2 diabetes and asthma. The approach captures clinically established mechanistic profiles in type 2 diabetes and asthma without requiring any reference phenotype data and enables identification of plasma proteins for novel subtype-specific biomarkers and drug targets. Lastly, we demonstrate that variants with conflicting effects on disease-relevant traits are resolved into distinct clusters, identifying subtypes with opposing phenotypic profiles that are lost by the aggregated genetic risk. Co-occurrence-based clustering provides a scalable strategy for understanding disease heterogeneity across diverse populations by identifying biologically meaningful subtype structure from genotype data alone.

## Introduction

Disease heterogeneity has been demonstrated across diverse diseases, marked by distinct risk factors, progression, response to treatment and biomarker expression^1–5^. Existing subtyping methods largely rely on disease-relevant clinical measures to delineate groups of individuals with shared characteristics^6,7^. However, clinical subtyping does not guarantee meaningful mechanistic differences between subtypes, which is crucial for both understanding disease outcomes and appropriate management^8,9^.

Genetic heterogeneity between clinically defined subtypes is considered to provide evidence of causal mechanistic differences^10^. This has led to the development of methods that directly leverage genetic data to understand disease heterogeneity^11–14^. The most widely used are partitioned polygenic risk scores (PGS), which partition an individual’s overall genetic risk by functional annotations^15,16^. Partitioned PGS have been shown to differentiate disease subtypes, but are limited by the accuracy and disease relevance of the reference annotations^11^.

To overcome these limitations, studies have examined how disease associated variants are associated with diverse traits to define variant clusters. For example, this approach has been successfully applied to Type 2 Diabetes (T2D) by Smith *et al*.^14^ and Suzuki *et al.*^13^ T2D is a metabolic disorder characterised by chronic hyperglycaemia, caused by inadequate insulin secretion from pancreatic beta-cells and increased insulin resistance in peripheral tissues^17,18^. Smith *et al*.^14^ and Suzuki *et al.*^13^ clustered T2D associated variants using associations with glucose homeostasis, liver function, adiposity and other T2D relevant traits to identify variant sets associated with key mechanistic processes with distinct phenotypic outcomes. The disease burden of T2D is largely attributable to its comorbidities; cardiovascular disease, chronic kidney disease and metabolic dysfunction associated liver disease. Given that T2D is a result of complex interactions between genetic and environmental risk factors, disease trajectories between individuals are highly heterogeneous^17,19^. Current clinical strategies cannot accurately predict patient outcomes, creating a need for alternative strategies to understand the mechanistic basis of disease to guide appropriate management.

However, studies like Smith *et al.*^14^ require knowledge of disease relevant phenotypes and high-quality phenotypic data capturing these traits at appropriate time points. For many diseases relevant endophenotypes^20^ are unknown. Moreover even if they are known, diseases are commonly characterised by transient, tissue-specific processes^21,22^, making it difficult to capture these processes because they demand time-sensitive, scalable, and complex sampling^23,24^. These challenges are further compounded in underrepresented populations, where ancestry-specific clinical features are poorly characterised.

To address these limitations, emerging approaches enable analysis of disease heterogeneity directly from genetic data without reliance on robust phenotypic reference points. One such method, BUHMBOX, leverages the principle that a disease may include a subtype that is genetically more similar to a secondary trait/disease, hence variants associated with a secondary trait that will be positively correlated in a heterogeneous cohort^25^. However, BUHMBOX requires prior knowledge of both a subtype-relevant secondary trait, and the variants associated with it, restricting its application in poorly characterised diseases^25^. Moreover, it only tests for evidence of a subtype, not which individuals are allocated to the subtype. Nonetheless, BUHMBOX provides evidence that when a subtype is present, variants associated with the subtype will co-occur more frequently than expected by chance.

There is a pressing need for a generalisable approach to uncover subtype-specific disease mechanisms from genetic data without requiring prior knowledge of the relevant traits or pathways. This study develops a phenotype-agnostic framework to identify genetic variants associated with disease subtypes by quantifying variant cooccurrence across individuals with the same diagnosis. Applied to simulated heterogeneous diseases and to T2D and asthma our method uncovers mechanistically distinct variant clusters that stratify individual disease outcomes using only genetic data. These subtypes map to both known and novel biomarkers, enabling clinically actionable insights from genetic findings. This generalisable approach enables unbiased discovery of disease mechanism without reliance on clinical data.

## Results

We hypothesized that in a disease cohort comprised of unknown subtypes with genetic differences, variants associated with the same subtype would co-occur across individuals more frequently. Quantifying the variant co-occurrence would therefore provide a phenotype free approach to identify sets of genetic variants that are causally and mechanistically associated with disease subtypes **(Figure 1A)**. Therefore, using variant co-occurrence to derive partitioned PGS could enable unsupervised stratification of individuals by their risk across subtype-specific pathways and inform novel biology of disease heterogeneity **(Figure 1C)**.

**Figure 1.**
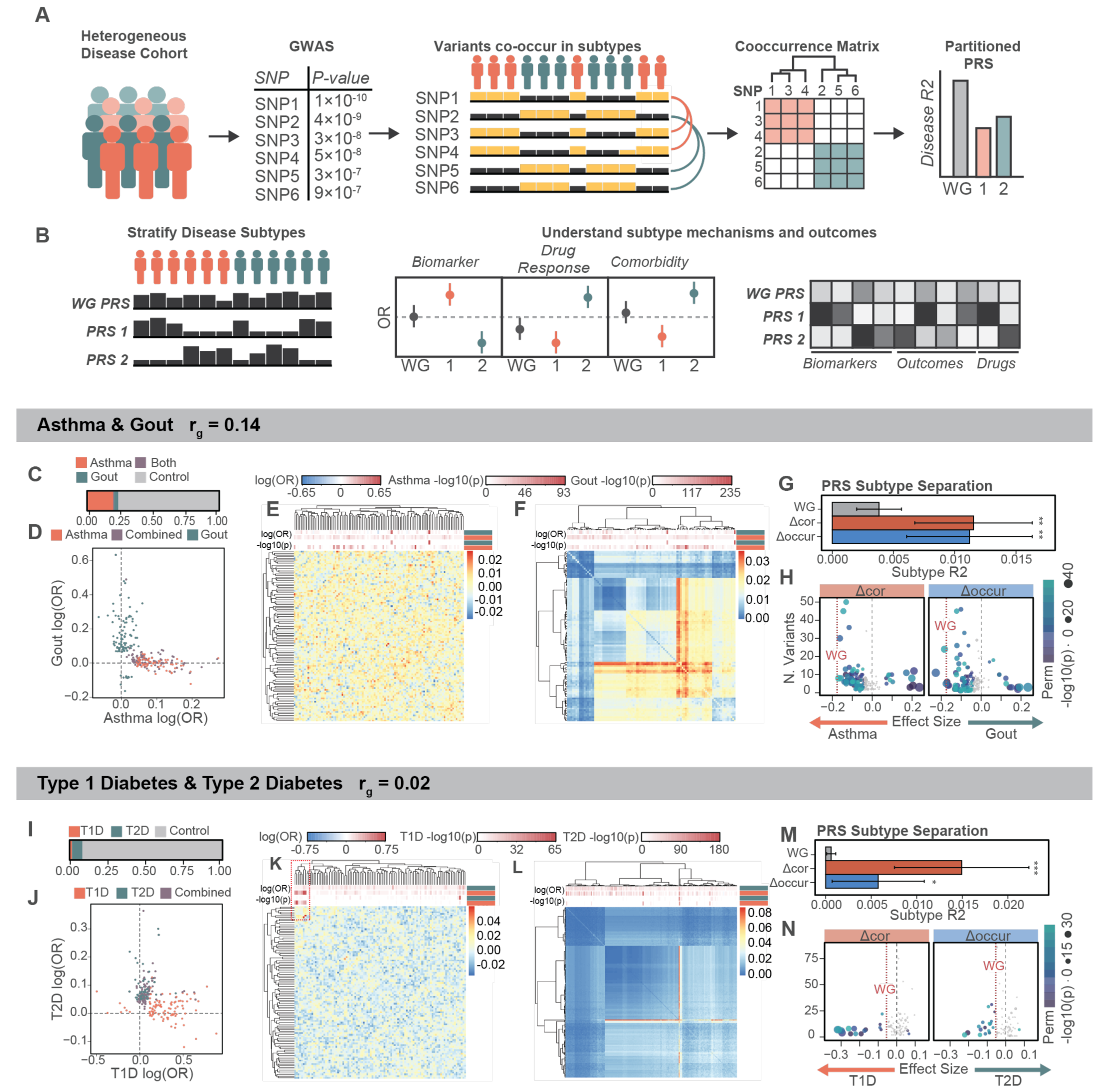
Rationale and validation of variant-cooccurrence to identify variant sets underlying disease heterogeneity. **(A)** Variants associated with a heterogeneous disease cohort will have distinct, quantifiable patterns of cooccurrence across individuals with different subtypes. Variant cooccurrence can be used to define variant sets and calculate partitioned PGS. **(B)** Partitioned PGS can stratify individuals into subtypes and reveal the mechanistic basis of subtypes by their association with diverse phenotypes and biomarkers. **(C-D)** Composition **(C)** and GWAS associations of composite disease phenotype made from individuals with asthma and/or gout. **(E-F)** Example **(E)** Δcor and **(F)** Δoccur matrices annotated by the effect size and significance in separate asthma and gout GWAS. **(G)** Proportion of subtype variance (Nagelkerke’s R^2^) explained by jointly fitting all cluster-partitioned PGS compared to the unpartitioned PGS. **(H)** Effect size of each cluster-PGS in separating asthma from gout across 10 permutations. **(I-N)** Analyses repeated for a composite cohort of T1D and T2D as per (C-H). Error bars indicate standard deviation. p < 0.05 * p < 0.01 ** p < 0.001 ***. WG: Whole Genome.

In this study, we apply two models to quantify variant-variant co-occurrence.

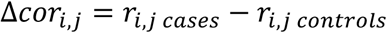

Δcor quantifies the difference in the correlation between disease associated variants *i* and *j* in the case and control cohorts respectively. The variants are those considered independent in the population (e.g. through stringent variant clumping/pruning given the correlation (linkage disequilibrium, LD) between them). Variants with increased correlation in case compared to controls are likely to be associated with the same disease subtype. However, in the presence of complex LD and population structure (which generates LD across chromosomes^26^), Δcor may be prone to detect outlier values that contribute to unstable cluster assignments. To mitigate against this, we also define Δ 𝑜𝑐𝑐𝑢𝑟⍰𝑖, 𝑗⍰ as

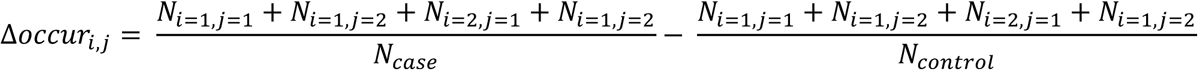

Δoccur quantifies the number of individuals with risk genotypes, defined as carrying either one or two copies of the risk allele, for disease associated variants *i* and *j* in the case and control cohort respectively. As a result, Δoccur captures increased cooccurrence of disease-associated variants in the case cohort that arises from increased frequency of risk alleles in affected individuals compared to control individuals. In the presence of confounders and complex genetic architecture, this signal may help to improve clustering stability and reveal the variants underlying subtype differences.

### Stratifying Individuals in Real Composite Phenotypes

To evaluate the performance of each model, we simulated heterogeneous disease cohorts by combining individuals with distinct diagnoses into a single cohort. Each composite phenotype cohort was partitioned equally into a training and testing set. Within the training cohort, a GWAS was conducted to identify the top 100 independent variants (r^2^ < 0.01) associated with the composite disease and estimate effect sizes. We then quantified Δcor and Δoccur and performed hierarchical clustering to define *n* variant clusters **(Figure S1A)**. Variant clusters were used to calculate *n* partitioned PGS in the testing set. The PGS used SNP weights from the composite phenotype GWAS. In the training set, the *n* PGS were jointly fit against the known subtype status (pre-composite disease label) to evaluate the proportion of subtype variance explained compared to an unpartitioned PGS. This analysis was repeated across 10 training-testing cohort permutations.

We tested this analysis in a composite cohort of individuals with either asthma (N=49,914) or gout (N=9,882) and 193,430 controls who had neither condition and met inclusion criteria to serve as controls for both traits. The genetic correlation (rg) estimate by LDSC regression was rg = 0.14, p = 8.6e-05 **(Figure 1C).** The 100 most associated SNPs from the composite trait were enriched for variants with shared effects on both traits and were completely depleted of variants with opposing directions of effect **(Figure 1D)**. Both Δcor and Δoccur showed stratification of variants with differing associations to asthma and gout **(Figure 1E-F)**. Furthermore, the Δcor and Δoccur derived variant clusters significantly outperformed the unpartitioned PGS in distinguishing individuals with gout from individuals with asthma across 10 permutations (p < 0.01) **(Figure 1G)**. Both approaches were consistently able to identify clusters with opposing directions of effect in separating each disease, suggesting clusters were capturing disease-specific genetic signals **(Figure 1H).** Importantly, the disease-specific signal was robust to the effect sizes used to calculate the PGS, maintaining direction of effect regardless of whether the phenotype weights were derived from asthma, gout, or a composite **(Figure S1B).** These clusters also reflected known biology: for example, a cluster consistently associated with increased risk of gout contained rs4148155, which is linked to *ABCG2*, a key gout-associated gene involved in urate transport^27^.

We next tested our framework in a composite phenotype of type 1 diabetes (T1D) (N= 3,136) and type 2 diabetes (T2D) (N=23,253) and controls (N=312,518) (rg = 0.02, p = 0.26) **(Figure 1I)**. The genetic architecture of these traits is highly distinct, where insulin-dependent T1D diabetes is driven by a few high effect variants largely concentrated in the HLA region^28^ **(Figure 1J)**. There was clear stratification of variants associated with T1D from those associated with T2D in Δcor, and lesser separation in the Δoccur **(Figure 1K-L).** While both approaches significantly outperformed the unpartitioned PGS (p < 0.05), the Δcor derived clusters explained more variance than Δoccur derived clusters **(Figure 1M)**. This gain in performance arose from Δcor consistently identifying clusters consisting of variants that are strongly associated with T1D **(Figure 1N, Figure S1C).**

We further validated our framework across composite cohorts comprised of individuals with asthma and chronic obstructive pulmonary disease (COPD) (rg = 0.66, p = 3.8×10^-148^), asthma and hypertension (rg = 0.162, p = 2.9e-11) and hypertension and heart failure (rg = 0.75, p = 1.9e-123) **(Figure S2A-C)**. Both Δcor and Δoccur derived clusters explained more subtype variance for asthma and COPD and asthma and hypertension than the unpartitioned PGS (p < 0.01) and captured clusters with opposing directions of effect **(Figure S2D-E)**. Interestingly, neither the unpartitioned PGS, nor either model could stratify individuals with heart failure from individuals with hypertension, likely due to hypertension acting as a major risk factor for heart failure, rather than as a distinct disease (consistent with the high rg) **(Figure S2F)**.

Finally, we investigated how these methods performed when the number of variants used to partition disease subtypes was increased from 100 to 250, 500, 1000 and 1500. With the exception of T1D and T2D, Δoccur derived clusters performed significantly better than the unpartitioned PGS, while Δcor quickly lost accuracy with increasing SNPs selected for the PGS **(Figure S2D-L)**. Worse performance at higher numbers of variants may be attributed to high clustering instability in Δcor, as indicated by the shallow dendrograms **(Figure 1E)**.

### Partitioning disease subtypes in type 2 diabetes

To evaluate the utility of quantifying variant co-occurrence in a real disease with known heterogeneity, we applied our framework to T2D. We sought to determine whether variant cooccurrence could recapture the mechanistic clusters identified by Smith *et al*.^14^ without requiring reference phenotypes. We first quantified Δcor and Δoccur for 650 independent T2D associated variants identified by Smith *et al*.^14^ in a cohort of 23,353 T2D cases and 312,518 controls in the UK biobank. While the 650 T2D- associated variants were selected from a GWAS that includes the UK Biobank, we did not test for association between these variants and T2D status, minimising bias. We used hierarchical clustering to derive 12 clusters from Δcor and Δoccur to match the number of clusters used by Smith *et al*.^14^ **(Figure 2A-D)**. Notably, Δcor-derived clusters were highly unstable, with each cluster conserved in only 14.4% of permutations when 15% of the T2D cohort was removed during the calculation of Δcor **(Figure S3A)**. In contrast, the Δoccur derived clusters were significantly conserved in an average 57% of permutations **(Figure S3B)**.

**Figure 2.**
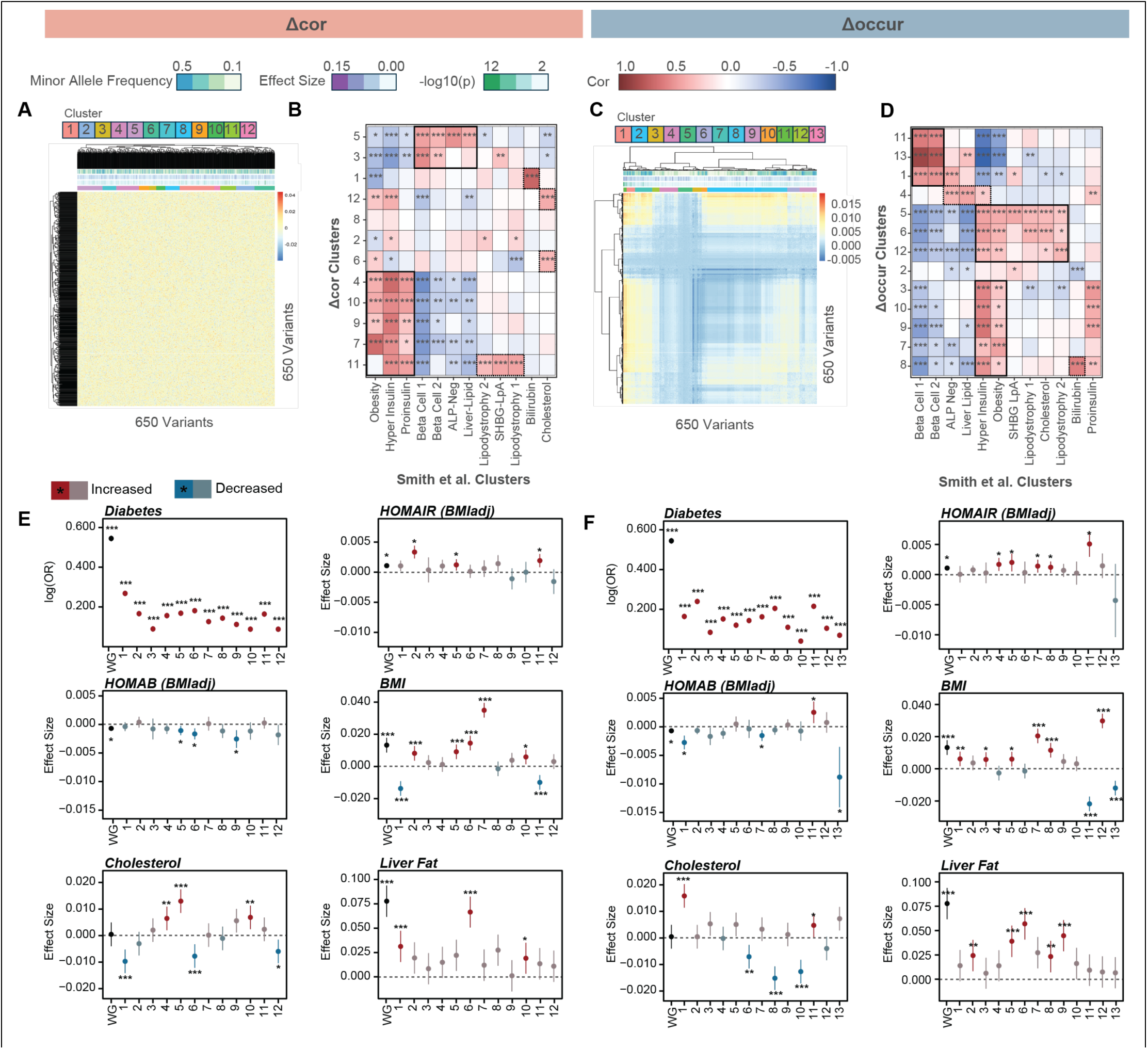
Variant cooccurrence recaptures phenotype-based mechanistic clusters in T2D. **(A)** Variant-variant correlation matrix of 650 T2D-associated variants divided annotated by effect size, minor allele frequency, p-value and cluster assignment. **(B)** Correlation between Δcor clusters and clusters identified by Smith *et al*. (2024) by aggregated effect sizes across 123 summary statistics. **(C-D)** As per (A-B) using for Δoccur. **(E-F)** Association of cluster-partitioned PGS and AES to key T2D risk factors and traits for **(E)** Δcor and **(F)** Δoccur derived clusters. Effect size shows change in standard deviation of trait per change in standard deviation of PGS or AES. Error bars indicate 95% confidence intervals. p < 0.05 * p < 0.01 ** p < 0.001 ***.

### Recapturing functional profiles identified by Smith et al

To functionally profile the clusters, we extracted variant–trait effect sizes from publicly available GWAS summary statistics across 123 curated phenotypes, broadly spanning measurements of glucose homeostasis, blood biomarkers, lipid levels, and body composition. For each cluster, we then performed inverse-variance weighted meta-analysis to compute aggregated effect sizes (AES), reflecting the overall direction and magnitude of each cluster’s association with each trait **(Supplementary Table 1)**. We evaluated the correlation in AES between the Δcor and Δoccur derived clusters and the clusters identified by Smith *et al.*^14^ to determine if similar mechanistic profiles were uncovered. Broadly, Δcor derived clusters partitioned into three groups; those related to insulin deficiency and pancreatic dysfunction, obesity and insulin resistance, and other more specific mechanisms, such as cholesterol metabolism **(Figure 2B)**. Similarly, Δoccur derived clusters partitioned into four groups: insulin deficiency, insulin resistance, specific mechanisms, and an additional group capturing lipodystrophy related clusters **(Figure 2D)**. While these groups are broadly consistent, there is considerable overlap between insulin-resistance and lipodystrophy clusters, reflecting shared metabolic features. Clusters from Δcor and Δoccur that shared functional profiles with clusters from Smith *et al*.^14^ also exhibited enrichment for variant overlap between them, though there was limited statistical significance **(Figure S3C-D)**.

### Functional profiles of correlation and cooccurrence derived clusters

Although all Δcor and Δoccur derived clusters were significantly associated with increased risk of T2D, they showed distinct signatures of association in both AES and partitioned PGS analysis **(Figure 2E-F, Supplementary Table 2)**. The associations of Δcor and Δoccur clusters across key T2D traits further reinforced shared biological mechanisms with clusters from Smith *et al*.^14^. For instance, clusters positively correlated with insulin deficiency-related clusters from Smith *et al*.^14^ were also negatively associated with HOMA-β, a metric of adequate insulin production. Clusters positively correlated with insulin resistance clusters from Smith *et al*.^14^ showed positive associations with BMI, waist to hip ratio, liver fat and HOMA-IR, a measure of insulin resistance **(Figure 2E-F, Figure S3E-F)**. Notably, cooccurrence-based clusters showed opposing directions of effect on BMI, blood cholesterol, apolipoprotein B, low-density lipoprotein, and mean corpuscular volume **(Figure 2E–F, Figure S3E-F)**, suggesting that variant co-occurrence effectively partitions T2D associated variants with conflicting effects on these traits. Indeed, the unpartitioned PGS had a weak, or no association with these traits, highlighting how aggregating variants with conflicting effects across disease subtypes can obscure associations with clinically meaningful risk factors.

To identify subtype-specific biomarkers and candidate drug targets, we calculated AES for each cluster using summary statistics for 2,923 plasma proteins in the UK Biobank^29^. Each cluster showed distinct, highly specific protein associations that were obscured by the aggregated genetic effects of all variants **(Figure S3G-H)**. Of particular note, Δoccur cluster 4 was associated with decreased PCSK9, while the aggregated genetic risk and obesity-related clusters are associated with increased PCSK9. PCSK9 inhibitors are used to manage the risk of cardiovascular disease by decreasing low-density cholesterol, however clinical studies have found conflicting evidence on whether PCSK9 inhibition increases the risk of developing T2D^30^. The opposing associations of PCSK9 to distinct T2D variant clusters suggest that the effect of PCSK9 inhibition may be context-dependent on the underlying mechanism of T2D risk.

We next investigated the associations of specific clusters across diverse clinical traits and biomarkers to explore the underlying mechanism. Both the PGS and AES for Δoccur cluster 7 were more positively associated with increased BMI and increased relative body size at age 10 than the unpartitioned PGS or other clusters **(Figure 3A)**. Furthermore, this cluster was non-significantly associated with decreased visceral fat after adjusting for BMI, and with a reduced ratio of visceral to subcutaneous fat compared to the unpartitioned PGS and other clusters **(Figure 3A)**. Together, this suggests a non-lipodystrophic obesity phenotype, marked by increased body weight from childhood. Indeed, plasma protein analysis showed negative associations with both ADIPOQ and SHBG, which have been implicated in metabolic syndrome and childhood obesity^31^ **(Figure 3B)**. Lastly, partitioned genetic correlation analysis showed that phenotypes associated with cluster 7 were more strongly genetically correlated with each other when using only cluster 7 variants, compared to using all T2D-associated variants **(Figure 3C).** Together, this suggests shared genetic regulation of these traits by the variants in cluster 7 that is not shared across all T2D variants. Similar findings were observed for Δoccur clusters 1 and 9, which were associated with phenotypes marking a profile of elevated cardiovascular risk and visceral adiposity respectively **(Figure S4A-F)**.

**Figure 3.**
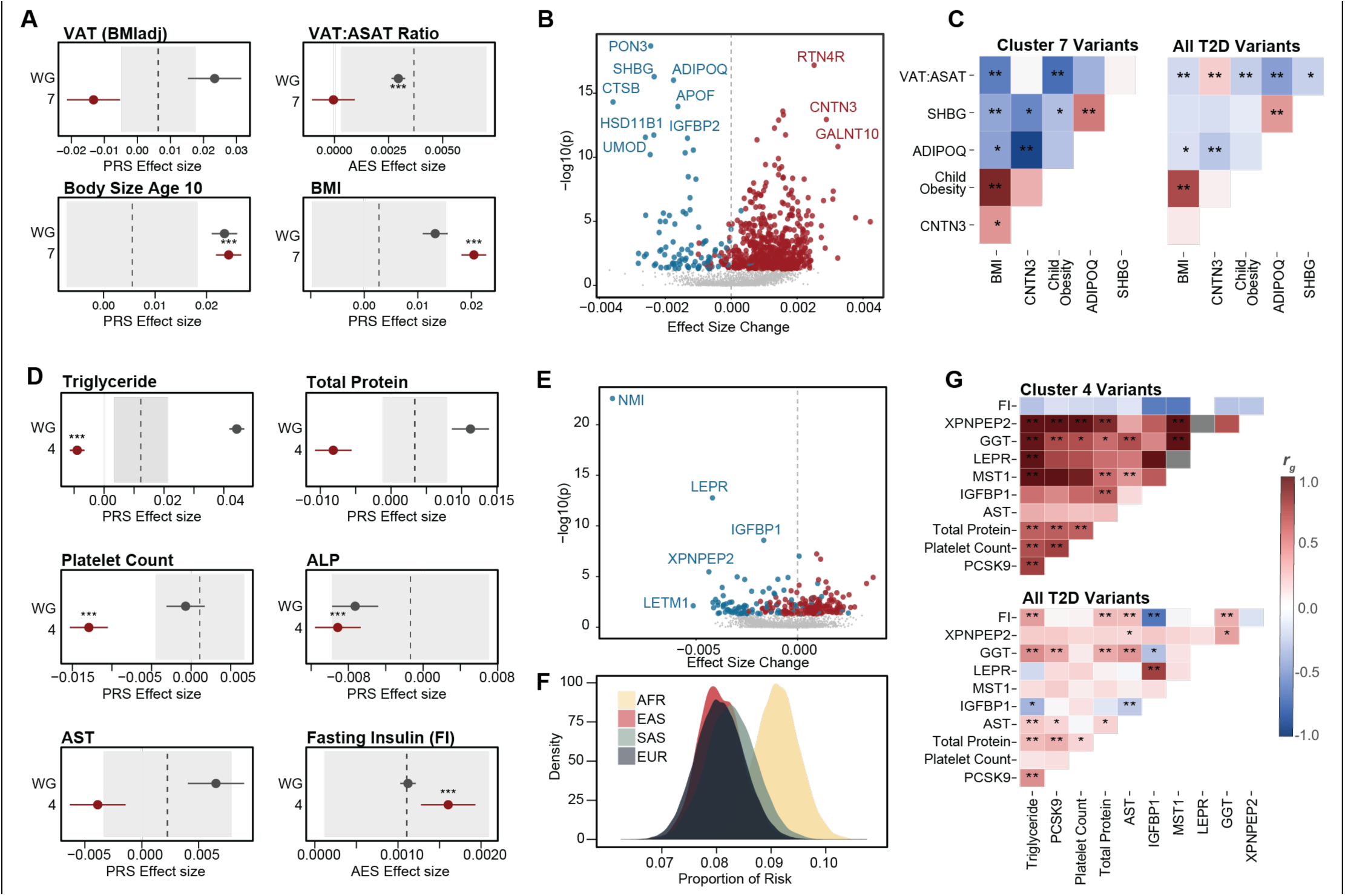
Cooccurrence derived clusters show shared genetic regulation of coherent mechanistic profiles. **(A)** Associations for distinguishing traits for Δoccur cluster 7 (AES and PGS). The aggregated variant set (WG) is provided as reference. Dashed line with grey box shows the average ± standard deviation of all other cluster eCect sizes. **(B)** Distinguishing plasma protein associations of cluster 7 compared to all variants. **(C)** Partitioned genetic correlation analysis between cluster 7 associated traits and biomarkers, partitioned by cluster 7 variants or all T2D variants. **(D)** Distinguishing traits for Δoccur cluster 4. **(E)** Distinguishing plasma protein associations for cluster 4. **(F)** Proportion of overall T2D risk attributable to variants in cluster 4 across diCerent ethnicities. **(G)** Partitioned genetic correlation analysis between cluster-4 associated traits and biomarkers, partitioned by cluster 4 variants or all T2D variants. p < 0.05 * p < 0.01 ** p < 0.001 ***

Lastly, we highlight Δoccur cluster 4, which appears to correspond to reduced hepatic clearance of insulin, which has been proposed as a primary cause of T2D^32^. Cluster 4 PGS was negatively associated with markers of liver function, including total protein in blood, platelet count, liver enzymes alkaline phosphatase and aspartate transferase, and blood triglycerides **(Figure 3D)**. Furthermore, the cluster 4 AES is associated with increased fasting insulin. Biomarker analysis shows decreased IGFBP1 and LEPR (leptin receptor), which have both been implicated in mediating liver sensitivity to insulin^33,34^ **(Figure 3E)**. Cluster 4 was also found to carry excess disease risk in the African population in the UKB, where reduced hepatic clearance of insulin and hyperinsulinemia have been shown to be more strongly associated with T2D risk than in European populations^32,35^ **(Figure 3F)**. Finally, we show that the biomarkers and phenotypes associated with cluster 4 were strongly genetically correlated when partitioned by the cluster 4 variants as opposed to all T2D variants, suggesting shared regulation of these processes **(Figure 3G)**.

### Predictive performance in T2D

Cluster partitioned PGS improve the prediction of individual disease risk for T2D and across broad comorbidities. We first tested whether cluster-partitioned PGS could identify diabetic individuals with low overall genetic risk for T2D but elevated risk within a specific disease pathway. Indeed, diabetic individuals with low total T2D PGS were significantly overrepresented compared to non-diabetic individuals among those with high cluster-specific risk, supporting the idea that concentrated genetic burden in a single pathway can drive disease (OR > 1.14, p < 6 × 10⁻⁸) **(Figure 4A-B)**. Partitioning variants using Δoccur and Δcor improved prediction of T2D related traits and risk factors by capturing the biologically distinct processes that contribute to these phenotypes. Both Δoccur and Δcor captured significantly more variance than the unpartitioned PGS and tissue-partitioned PGS in predicting T2D- related traits and risk factors. Notably, the greatest performance gains occurred for traits with opposing directional effects across clusters, such as Apolipoprotein B **(Figure 4C)**.

**Figure 4.**
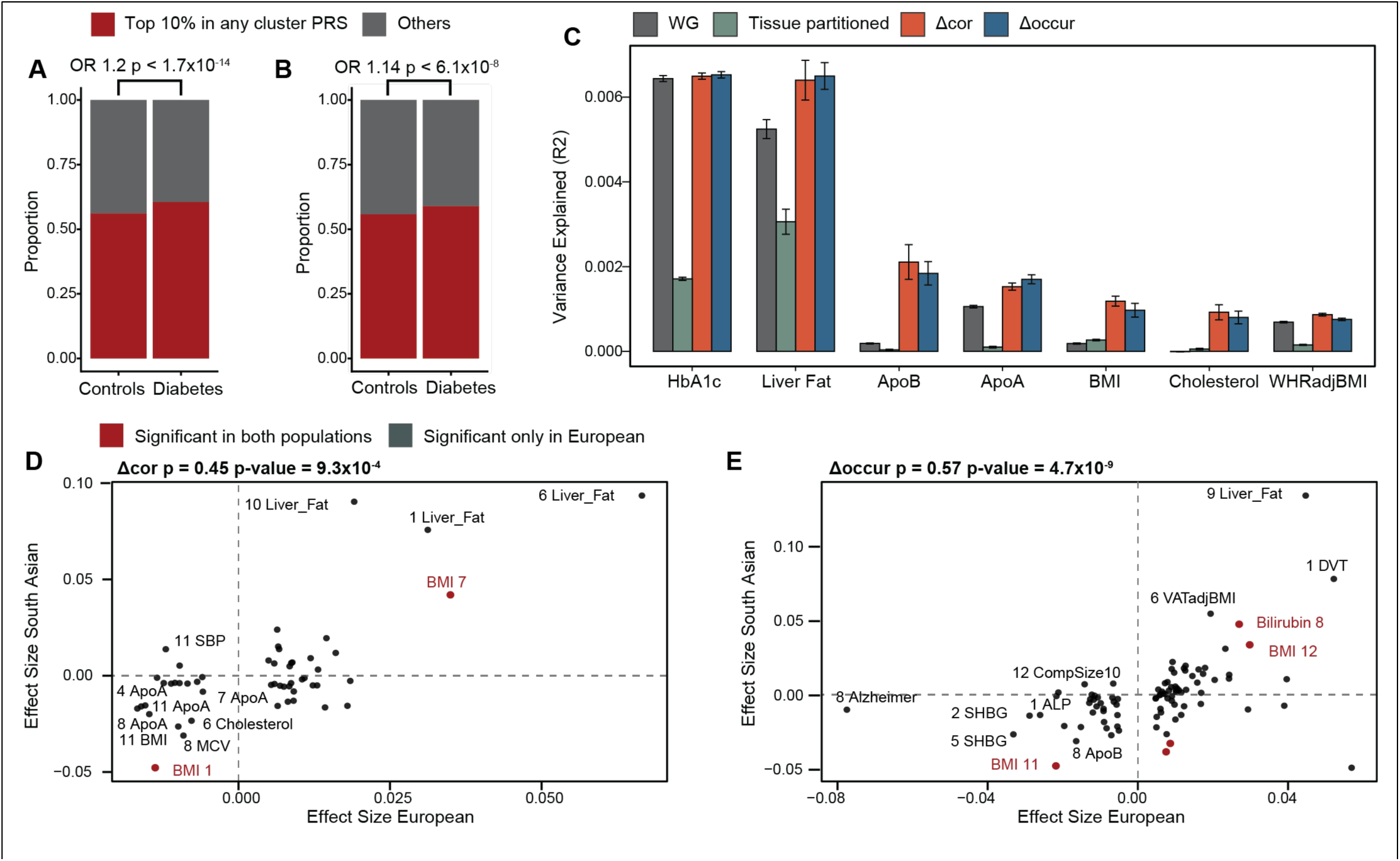
Validation and application of cooccurrence derived clusters in T2D. **(A-B)** Individuals with T2D but low overall genetic risk for T2D risk are detected in the top 10% of at least one cluster-partitioned PGS in both **(A)** correlation and **(B)** cooccurrence. Chi-squared enrichment test reported. **(C)** Proportion of phenotypic variance (R2) explained by jointly fitting all cluster partitioned-PGS compared to the unpartitioned PGS and tissue-partitioned PGS across 10 permutations. **(D-E)** Correlation of eCect sizes for each cluster-PGS and phenotype pair between European and South Asian cohorts for **(D)** Δcor and **(E)** Δoccur derived clusters. Significant associations in the South Asian cohort (p < 0.05) are highlighted. Spearman’s correlation coeCicient reported. Error bars show standard error. p < 0.05 * p < 0.01 ** p < 0.001 ***

### Generalisability of clusters across ancestry

To determine if variant clusters derived using co-occurrence across individuals were confounded by population structure, we evaluated the correlation between cluster-partitioned PGS and the first 10 genetic components. We found minimal correlations with the first 10 genetic principal components **(Supplementary Table 3)**. Furthermore, the direction and magnitude of cluster-specific effects observed in the Caucasian population were broadly conserved in the South Asian population (ρ > 0.45, p < 0.001), despite a limited cohort size in the UKBB **(Figure 4D-E)**. Importantly, this included traits like BMI where opposing directions of effect were identified in both Caucasian and South Asian populations, reinforcing their biological robustness.

These findings were replicated for the T2D associated loci and clusters identified by Suzuki *et al*.^13^ (Figure S5, Figure S6, Supplementary Tables 4-5).

### Applying variant cooccurrence to identify mechanistic clusters in asthma

To evaluate the broader utility of our framework, we applied it to asthma, a trait with clinical heterogeneity that lacks established genetic subtypes. Asthma is a complex, immune-mediated disease characterised by airway inflammation^20^. Clinical studies have identified subtypes involving eosinophilic inflammation, neutrophilic inflammation and obesity-related asthma that each have differences in patient biomarkers and characteristics^20^. However, robust molecular subtyping of asthma is challenging due to both spatially and temporally restricted inflammatory processes^36,37^. Our framework provides a unique opportunity to dissect heterogeneity in asthma without requiring in depth clinical data.

We applied our framework to 116 independent (r^2^ < 0.01), genome-wide significant variants associated with asthma to identify variant clusters underpinning molecular heterogeneity^38^. We evaluated cluster numbers from 2 to 15 for each method and performed IVW meta-analysis across a panel of 93 phenotypes **(Supplementary Table 6)**. Splitting the Δcor and Δoccur matrices into 8 clusters provided the greatest separation between clusters in their functional profiles while maintaining sufficient variants for statistical power **(Figure S7A-D)**. As in T2D, the Δcor matrix was susceptible to minor perturbations in the case cohort, while the Δoccur matrix was more robust **(Figure S7E-G)**.

We calculated partitioned PGS for each cluster and tested the associations between the PGS and a panel of asthma related phenotypes both in the full UKBB cohort adjusting for asthma status, and in asthmatic individuals only. Though nearly all clusters were positively associated with asthma risk, they showed distinct patterns of association across key asthma biomarkers **(Figure 5A).** For instance, both Δcor and Δoccur identified two highly eosinophilic clusters, associated with higher prevalence of hay fever, high eosinophil percentage, low neutrophil percentage, and early onset of asthma, relative to other clusters **(Figure 5A)**. Furthermore, these clusters had strong, positive associations to plasma proteins associated with eosinophil activation and degranulation, such as PRG2/3 and IL5RA, when compared to clusters with a non-eosinophilic profile **(Figure 5B)**. Similarly, we were able to identify clusters associated with neutrophilic asthma, marked by an increase in both neutrophil counts and percentage, and reinforced by plasma protein associations **(Figure 5A & C)**. We also identified obesity-related asthma clusters, characterised by higher BMI and later asthma onset **(Figure 5A).** Notably, traits such as BMI and neutrophil percentage were not associated with the unpartitioned PGS, but showed divergent associations across clusters, suggesting that clustering disentangles opposing effects that are otherwise masked in aggregate genetic risk. These cluster-phenotype signatures were broadly conserved in the South Asian cohort in the UKBB **(Figure S8A-B).** Furthermore, asthmatic individuals with low total asthma PGS were significantly overrepresented among those with high cluster-specific risk, supporting the presence of concentrated, pathway-specific genetic burden **(Figure S8C-D)**. Lastly, we show that jointly fitting partitioned PGS against key asthma traits captured more variance than the unpartitioned PGS in both the full cohort and the asthma cohort **(Figure S8E-F)**.

**Figure 5.**
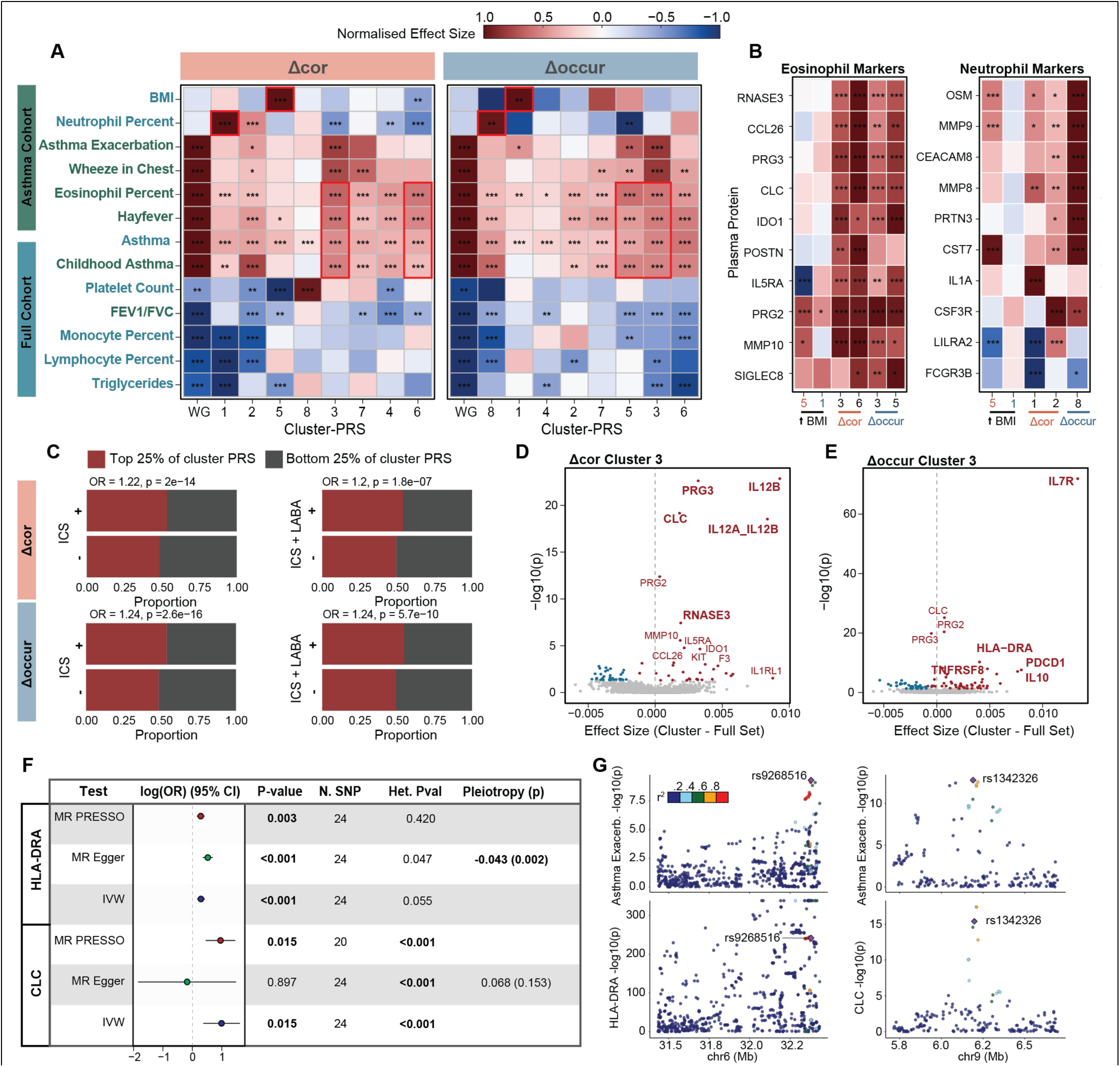
Variant cooccurrence identifies clusters with distinct mechanistic profiles in asthma. **(A)** Cluster partitioned PGS associations across a panel of asthma-related traits in the full cohort and in asthma-only cohort. ECect size normalised by the maximum absolute value per trait. **(B-C)** Association of known **(B)** eosinophil markers with eosinophilic asthma clusters and **(C)** neutrophil markers with neutrophilic asthma clusters compared to obesity-related asthma clusters. **(D-E)** Enrichment for individuals taking inhaled corticosteroids (ICS) or inhaled corticosteroids and long-acting beta-agonists (LABA) in the top 25% of the **(D)** Δcor cluster 3 and **(E)** Δoccur cluster 3 partitioned PGS compared to the bottom 25%. **(F-G)** Distinguishing plasma proteins for clusters associated with higher risk of asthma exacerbation. Plasma proteins for mendelian randomization are highlighted. **(H)** Significant associations from MR analysis of plasma protein abundance as exposure on asthma hospitalisation as outcome. **(G)** Colocalization of protein QTLs with asthma hospitalisation loci for MR significant plasma proteins. p < 0.05 * p < 0.01 ** p < 0.001 ***

Finally, we identified variant clusters using both Δcor and Δoccur that are associated with an increased incidence of asthma exacerbations. Interestingly, the clusters most strongly associated with asthma exacerbations by both methods (Δcor cluster 3, Δoccur cluster 3) were associated with elevated eosinophil percentage, possibly indicating worse control of asthma-related inflammation **(Figure 5A)**. Indeed, asthmatic individuals with high genetic risk in these clusters had increased rates of using inhaled corticosteroids (ICS) (OR > 1.22, p < 0.001), and increased rates of dual treatment with ICS and long-acting beta agonists (LABA) (OR > 1.2, p < 0.001), which are often prescribed in individuals with insufficient response to ICS **(Figure 5C)**. Notably, a high PGS in Δoccur cluster 3 was associated with significantly increased use of ICS monotherapy and combination ICS/LABA therapy, even after accounting for the overall genetic risk (p < 0.01).

We identified plasma proteins^29^ upregulated in exacerbation-associated clusters compared to all asthma associated variants **(Figure 5D–E),** including IL7R, previously linked to corticosteroid resistance^39^, and IL5RA, a therapeutic target of monoclonal antibody treatment for severe asthma^40^. To test for evidence that these proteins could be causally associated with asthma exacerbation, we performed mendelian randomization analysis using three methods (MR PRESSO^41^, inverse-variance weighted analysis (IVW), MR Egger^42^) between the top 5 plasma proteins from each cluster and a GWAS of asthma hospitalisation in a paediatric cohort^43^ **(Figure S9A)**. Increased plasma HLA-DRA was significantly associated with increased asthma hospitalisation across all three methods and increased plasma CLC was significantly associated in MR PRESSO and IVW analysis **(Figure 5F, Figure S9B-C)**. We find that these associations are robust to the exclusion of any one SNP, as shown by leave-one-out sensitivity analysis, and that a number of protein-quantitative trait loci (pQTLs) for both HLA-DRA and CLC were colocalised (PP > 0.75) with loci associated with asthma exacerbation **(Figure 5G, Figure S9D-E)**. Notably, a colocalised locus, rs1342326, is a trans-pQTL for CLC located within IL33, a key driver of eosinophil-based inflammation in asthma^44^ **(Figure 5G)**. CLC was previously used as part of a gene expression signature in sputum that was predictive of asthma exacerbations but had not been causally associated^45^. Similarly, while HLA-DRA is known to be involved in inflammation, it has not been previously linked to asthma exacerbation.

Together these findings highlight how our framework can identify genetic subtypes of disease with distinct mechanisms that can guide biomarker and novel drug target identification for clinical management, without requiring in-depth patient characterisation.

## Discussion

Partitioning genetic risk by subtype-relevant genetic pathways is a powerful strategy to identify individuals with distinct clinical outcomes and understand the mechanistic processes driving them. This study develops an unsupervised framework to reveal genetic pathways driving subtype differences using only genetic data. We show that quantifying the cooccurrence of variants in a heterogeneous cohort is sufficient to partition individuals into their true subtypes using composite phenotypes.

We applied our framework to T2D, which has been the focus of many clinical and genetic studies to understand patient heterogeneity, producing in-depth phenotypic data. We use T2D to demonstrate that variant cooccurrence can recapture mechanistic profiles identified by prior works using extensive phenotypic data^13,14^, finding biologically coherent clusters across cardiovascular disease, dyslipidaemia, liver dysfunction and adiposity-related traits. Even with an abundance of reference data, phenotype-based subtyping studies are likely to bias findings toward the phenotypes used to define variant clusters^10^. Quantifying the variant cooccurrence provides an unbiased, orthogonal approach to both validate discovered biological processes and identify novel disease clusters.

T2D is unusual among complex diseases, where its key mechanistic traits are both well-studied and measurable using routine clinical and biochemical data. Reference-based subtyping methods using curated phenotypic or functional annotations are directly constrained by the availability of high-quality reference data. For many diseases, such as asthma, capturing disease-relevant molecular data is challenging due to spatially or temporally restricted mechanisms. In these diseases, both clinical and molecular subtyping efforts have produced inconsistent results that are unstable to changes in the studied cohorts^37^. Clinical and genetic studies subtyping asthma rely on proxy phenotypes in place of direct molecular measurements, using blood eosinophilia as a measure of airway eosinophilic inflammation despite limited correlation between them^22,46^. The reliance on proxy phenotypes that fail to capture the underlying disease biology limits the accuracy and utility of subtyping efforts.

Our phenotype-free framework overcomes these limitations by inferring disease subtypes directly from genetic data. Applying our framework in asthma, we identify variant clusters with distinct inflammatory profiles that stratify individuals by clinically relevant characteristics, including a novel cluster associated with asthma exacerbation.

Although powerful to identify subtype-driven genetic structure, variant cooccurrence may be susceptible to detecting LD and subtle population structure, despite the use of independent variants and an ancestry matched control cohort. In particular, correlation derived clusters were sensitive to small changes in the cohort composition and prone to detect outliers. Large case cohort sizes are needed to robustly identify clusters using variant cooccurrence, which can reduce the accuracy of our framework in populations with limited genetic data. Finally, we note that although clusters are derived without external references, understanding the disease mechanism requires data linking variants to diverse phenotypes.

Quantifying variant cooccurrence provides a generalisable strategy to study disease heterogeneity in any complex disease with available genetic data. Compared to traditional analyses, our framework can resolve subtypes with opposing phenotypic profiles that are obscured by aggregated genetic risk and confound genetic studies. By uncovering subtype-specific associations between variant clusters and plasma proteins, we identify both established drug targets, such as IL5RA and novel candidates to improve diagnosis and personalise treatment. Lastly, this framework enables the discovery of population-specific risk factors that are obscured by lacking phenotypic data in underrepresented populations. Ultimately, we develop an approach to partition disease-associated variants into biologically meaningful pathways to support the development of subtype-specific PGS, biomarker panels for targeted disease diagnosis and management.

## Acknowledgements

This research has been conducted using the UK Biobank Resource under Application Number 91057. We acknowledge Judith Gonzalez Garcia for her guidance in designing the composite trait analyses.

## Author Contributions

**DM** and **NP** conceived the study. **DM** developed the methodology, performed the analyses and visualised the results. **DM** and **NP** drafted the manuscript. **NP, SS,** and **NW** supervised the project, provided statistical guidance and assisted in interpretation of results. All authors reviewed and approved the final manuscript.

## Methods

### UK Biobank Cohort and Quality control

The UKBB resource contains deep phenotypic and genetic data for approximately 500,000 individuals, all of whom provided informed consent^47^. Genotypes were imputed using the merged 1000 Genomes and UK10K reference panels released by the UKBB and were employed for all analyses.

Quality control (QC) was performed at both the individual and variant levels using PLINK (version 1.90) and PLINK2 (version 2.00) on the UKBB RAP (Research Analysis Platform), following the recommendations in Marees, et al. ^48^. Specifically, variants with genotype missingness > 1% and individuals with genotype missingness > 3% were excluded. Additionally, variants deviating from Hardy-Weinberg equilibrium (p-value < 1e-10) were removed. To control for relatedness, individuals with a kinship coefficient > 0.2 were identified, and one individual from each related pair was excluded.

For ancestry filtering, we retained individuals who self-identified as “White British” (Field 21000) and those with similar genetic ancestry (Field 22006) for the GWAS and variant co-occurrence calculations.

### Cohort definitions

Asthma was defined in the UKBB as per Han et al. (2020)^49^. Doctor diagnosed asthma, ICD-10 codes J45 and J46, and self-reported asthma were used to define asthmatic individuals. Non-asthmatic individuals were defined as those free from self-reported asthma, eczema, hay-fever and allergy, lacking doctor diagnosed allergic diseases, and free of ICD-10 codes J30 and L20, yielding 49,914 cases and 198,898 controls.

COPD was classified using lung function tests as described in Doiron et al. (2019)^50^. The FEV_1_/FVC ratio was calculated using the highest value for each, measured at the most recent time the participant attended the centre. To account for variations around age, sex and height, the Global Lung Function Initiative reference values were used to define the lower limit of normal for each participant. Individuals below the lower limit of normal were classified as COPD cases, yielding 26,323 cases and 246,804 controls.

Gout was classified as per Sandoval-Plata et al. (2021)^51^ yielding 9,882 cases and 331,976 controls. Individuals were included in the case cohort if they self-reported physician diagnosed gout, used urate-lowering treatment or had ICD-10 codes M10, M10.0, M10.1, M10.2, M10.3 M10.4 and M10.9. Individuals on urate-lowering treatment either without a diagnosis of gout or with a secondary diagnosis of leukemia (C90-96) or lymphoma (C81-C88) was excluded.

Heart failure was defined in the UKBB as described in Shah et al. (2020)^52^ yielding 12,278 cases and 329,145 controls. Individuals were defined as having heart failure if they self-reported ‘heart failure/pulmonary oedema’ or ‘cardiomyopathy’, or had an ICD-10 classification of heart/ventricular failure or cardiomyopathy (ICD-10 codes: I11.0, I13.0, I13.2, I25.5, I42.0, I42.5, I42.8, I42.9, I50.0, I50.1, I50.9; ICD-9 codes: 4254, 4280, 4281, 4289). Individuals who either self-reported or had an ICD- 10 based classification of hypertrophic cardiomyopathy were excluded.

Hypertension was defined as per Evangelou et al. (2018)^53^, yielding 173,566 cases and 150,832 controls. To define hypertensive individuals, the two systolic and diastolic blood pressure measurements for each participant were averaged. Those with an average systolic blood pressure ≥ 140 mmHg, diastolic blood pressure ≥ 90mmHg or those reported to take a blood pressure lowering medication were classified as hypertensive.

Individuals diagnosed with non-insulin-dependent diabetes mellitus (ICD10 code E11) were classified as T2D cases. Those who self-reported diabetes but did not have an E11 diagnosis or had ICD10 codes for Type 1 diabetes (E10) or other forms (E12–E14), or individuals taking glucose-lowering medication were excluded. Using these criteria, 23,353 individuals were classified as T2D cases and 312,518 as controls within the UKBB cohort. T1D cases were classified under similar criteria, with those matching the ICD10 code E10 classified as cases, and others excluded, yielding 3136 cases and 324,778 controls^54^.

### Variant selection

For each composite disease, the cohort in the UKBB was randomly split into a testing and training cohort across 10 permutations. Within each permutation, a GWAS of the composite phenotype was conducted in the training cohort, using age, sex, Townsend deprivation factor and the first 10 principal components as covariates. LD clumping was performed on the resulting summary statistics using the 1000 Genomes European genotypes. The top 1500 variants were selected for downstream analysis. To evaluate the effect of variant number on performance, the top 100, 250, 500, 1000 and 1500 variants were selected by their significance.

The 650 and 1,289 variants associated with Type 2 Diabetes (T2D) at genome-wide significance were used, as reported by Smith et al. and Suzuki et al., respectively. Variants with a MAF < 0.01 were mapped to a proxy variant with MAF > 0.01 that were in LD with the original index variants. Original index variants were used for all other analysis.

116 genome wide significant, independent loci (r < 0.01) with MAF > 0.01 were identified from the GBMI meta-analysis of asthma excluding the UKBB for downstream analysis.

### Variant Cooccurrence Calculation

Variant Δoccur and Δcor was calculated in half of the overall UKBB cohort. The effect allele was harmonised to the relevant summary statistic used so that all effect alleles increased the risk of disease. Any missing genotype values were imputed with the average value for the corresponding variant, rounded to the nearest whole digit.

### Cluster number optimisation

To obtain clusters, hierarchical clustering was performed on the Euclidean distance transformed cooccurrence and correlation matrices across k numbers of clusters, where 2 < k < 25.

To select the optimal number of clusters for the composite trait analysis, partitioned PGS were calculated for every value of k and jointly fit in the training cohort to stratify composite cases from controls. The first 10 principal components, sex, age and Townsend deprivation factor were used as covariates. The proportion of variance explained by the model was quantified for each value of k. A plateau spanning two continuous values of k was used to determine the optimal number of clusters for subtype separation.

The optimal number of clusters for diabetes were fixed at 12 and 9 for the Smith *et al*. and Suzuki *et al.* variant sets, respectively. To account for the high likelihood of outlier clusters in hierarchical clustering, we optimised the number of clusters so that 12 and 9 clusters had at least 10 variants each, for the two variant sets.

To identify the optimal number of clusters for asthma associated variants, we performed meta-analysis across 93 diverse traits profiling immune cell counts, allergy, anthropometric phenotypes, and common comorbidities of asthma. For each value of k, we evaluated the average, absolute, aggregated effect size, Jaccard correlation of clusters with the full variant set, inter-cluster Jaccard correlation, and the number of significant associations per cluster.

### Individual Subsampling

Across 100 permutations, 85% of the case cohort was randomly subsampled and variant correlation and cooccurrence calculations and clustering were performed as described above. The proportion out of 100 clustering iterations that two variants were assigned together was calculated for all variant-variant pairs. For each of the cooccurrence and correlation approaches, we evaluated the average proportion that a cluster’s variants were assigned together and performed 100-fold bootstrapping to obtain significance.

### Aggregation of cluster-variant effects across the phenome

We collated 6,346 summary statistics from the UKBB and external sources, spanning 1,245 phenotypes and 2,923 plasma protein measurements. These summary statistics were harmonized so that the effect allele always increased the risk of the disease. To aggregate the effect sizes of all variants in each cluster, fixed-effect inverse-variance weighted meta-analysis (IVW-MA) was performed, in alignment with the methodology in Suzuki et al. IVW-MA was performed for each phenotype-cluster combination and for all T2D or asthma associated variants. IVW-MA was also performed for the clusters derived by Smith et al. and Suzuki et al. FDR correction for multiple testing was performed accounting for the number of clusters and the number of phenotypes. 123 phenotypes were manually selected for T2D and 93 phenotypes for asthma to minimise redundancy and capture biological breadth and cross-cluster variability.

### Comparison of functional profiles

To compare functional profiles with the clusters established in Smith et al. and Suzuki et al. we calculated Pearson’s correlations between the aggregated effect sizes of cooccurrence derived clusters, and the clusters identified by Smith *et al*. and Suzuki *et al*. To account for large, shared effects on phenotypes highly correlated with T2D, aggregated effect sizes for each cluster were transformed to the deviation from the aggregated effect size of the full variant set. The deviation was nullified if the cluster-phenotype association was non-significant.

### Partitioned Correlation Analysis

Partitioned correlation analysis was conducted using LD score regression (https://github.com/bulik/ldsc). For the entire T2D variant set and for each cluster, a credible set of variants was defined within 1Mb and in high LD (r > 0.8) of each index SNP. These credible sets were used as annotations to estimate genetic correlations between selected panels of traits and biomarkers selected for each cluster. FDR correction for multiple testing was applied for the number of trait-pairs tested.

### PGS Calculation

Partitioned PGS were calculated for each cluster using a weighted sum of variants weighted by their effect size from either the GWAS conducted in the training cohort for the composite phenotypes or using external GWAS that did not include the UKBB (X *et al* for T2D and X *et al* for asthma). A PGS containing all studied variants was used as the whole-genome control. All PGS were standardized to have a mean of 0 and a standard deviation of 1.

### PGS Analyses

Cluster-PGS were tested for association with a panel of selected phenotypes in T2D and in asthma. Cluster-specific phenotypes were selected using the results from IVW-MA. Continuous phenotypes were standardised to have a mean of 0 and standard deviation of 1. With the exception of predicting T2D status, all PGS analyses for T2D were conducted in the full UKB cohort with adjustment for T2D status. PGS analyses for asthma were conducted in both the full UKB cohort with adjustment for asthma status, and in the asthma-only cohort. A logistic and linear regression was used for binary and continuous traits, respectively. The first 10 principal components, age, sex, Townsend deprivation index were used as covariates, and T2D/asthma medication status, lipid medication or hypertension medication status where required. The models were first fit in the training cohort for each PGS-phenotype pair, and if passing a nominal p-value threshold (p < 0.05) then applied in the testing cohort. FDR correction for multiple testing was applied for the number of PGS-phenotype pairs in the testing cohort.

### Proportion of variance captured by cluster-partitioned PGS

All clusters were jointly fit against the trait of interest to obtain the proportion of variance explained (R^2^). The proportion of variance explained by a null model fitting only covariates was used to estimate the gain in R^2^. The proportion of variance captured by a model fitting an unpartitioned PGS was used as reference.

Tissue-partitioned PGS were used as benchmark. We performed mapping of index variants to accessible chromatin regions in the CATLAS database as described by Smith *et al*. We selected the top 12 and 9 CATLAS tissues by their enrichment for T2D variants as benchmarking for the Smith *et al*. and Suzuki *et al.* analyses respectively. We calculated partitioned PGS for all SNPs accessible in each tissue and jointly fit them against subtype status.

To obtain error size estimates, this analysis was repeated across 10 permutations of training-testing cohorts with re-calculation of cooccurrence and correlation matrices.

### Ancestry Definition

Individual ancestry was defined based on the approach described in Sun, et al.^55^. The first four genetic principal components were used to perform K-means clustering on individuals, deriving four clusters. For each cluster, the proportion of individuals with each self-identified ethnicity was calculated, and majority groups were assigned. Individuals with self-identified ethnicity that did not align with the cluster’s most populous groups were excluded. This resulted in the following cluster-ethnicity match ups:

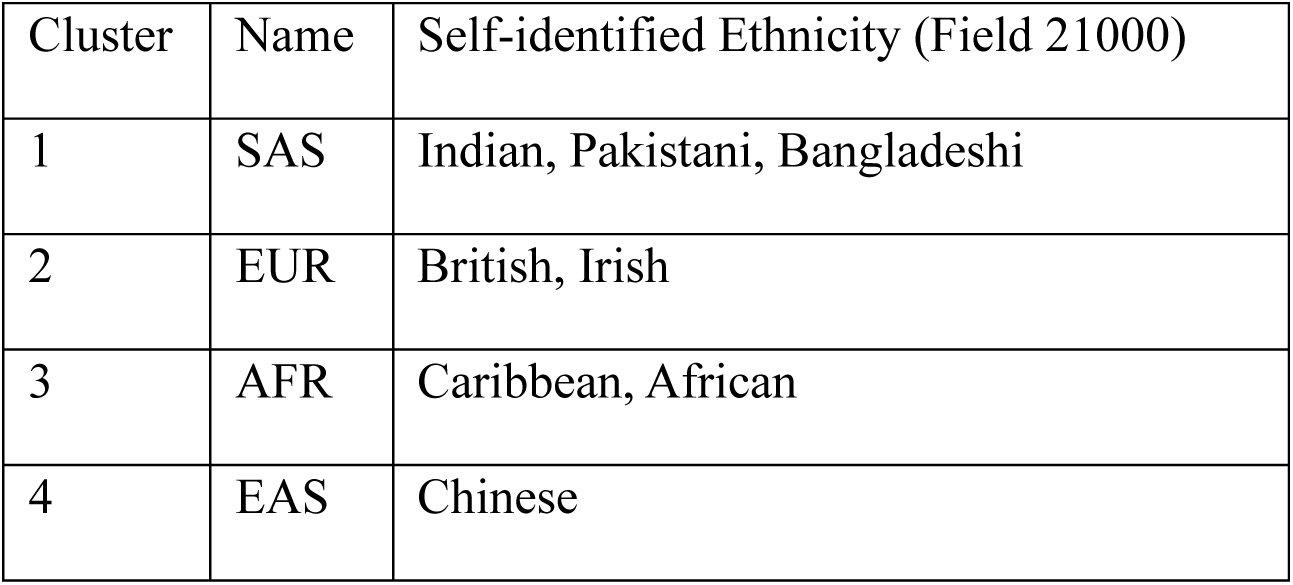

| Cluster | Name | Self-identified Ethnicity (Field 21000) |
| --- | --- | --- |
| 1 | SAS | Indian, Pakistani, Bangladeshi |
| 2 | EUR | British, Irish |
| 3 | AFR | Caribbean, African |
| 4 | EAS | Chinese |

Individual with ambiguous self-identified ethnicity (i.e. Any other Asian background, Other ethnic group) were included in the cluster, provided their Euclidean distance from the cluster centroid was < 100. This threshold was selected based on the distance distribution of individuals with self-identified ethnicities in alignment with the cluster ethnicity.

### PGS Replication in South Asian Cohort

All cluster-PGS associations that were significant in the European testing cohort were evaluated in the South Asian cohort. The full South Asian cohort, adjusting for T2D status or Asthma status was used. FDR correction for number of cluster-phenotype pairs was applied. To evaluate consistency of effect sizes between European and South Asian cohorts, spearman’s correlation coefficient was calculated using each cluster-phenotype effect size.

### Trans-ancestry Risk Burden

An individual’s total disease risk was defined as their cumulative risk across all cluster-PGS. To calculate cumulative risk, PGS were normalised using min-max normalisation. The proportion of the total risk attributable to each cluster was calculated for each individual as described in Smith et al., (2024). Individuals were stratified by their assigned ancestry to determine if specific pathways contributed more to disease risk in each population. A one-way ANOVA was used to determine statistical significance.

### Mendelian Randomization Analysis

We performed two-sample Mendelian Randomization to evaluate if the plasma proteins more highly associated with asthma-exacerbation clusters had a causal effect on asthma hospitalisation. Genome-wide significant, independent (r < 0.01) pQTLs were used as exposures. MR analyses were conducted using the *TwoSampleMR* (v0.5.7) and *MRPRESSO* (v1.0) R packages. Causal estimates were obtained using the IVW, MR-Egger and MR-PRESSO to account for presence of outlier SNPs or horizontal pleiotropy.

For CLC and HLA-DRA, leave-one-out analysis was performed to ensure robustness of the association. Further, colocalization analysis was performed using *Coloc* (v5.2.3) and visualised using *locuscompareR* (v1.0.0) R packages to test if the causal variant was shared between the protein exposures and asthma hospitalisation.

