## Supplemental Figures for "Unsupervised Variant Clustering Identifies Genetic Subtypes of Disease"

### Supplementary Materials

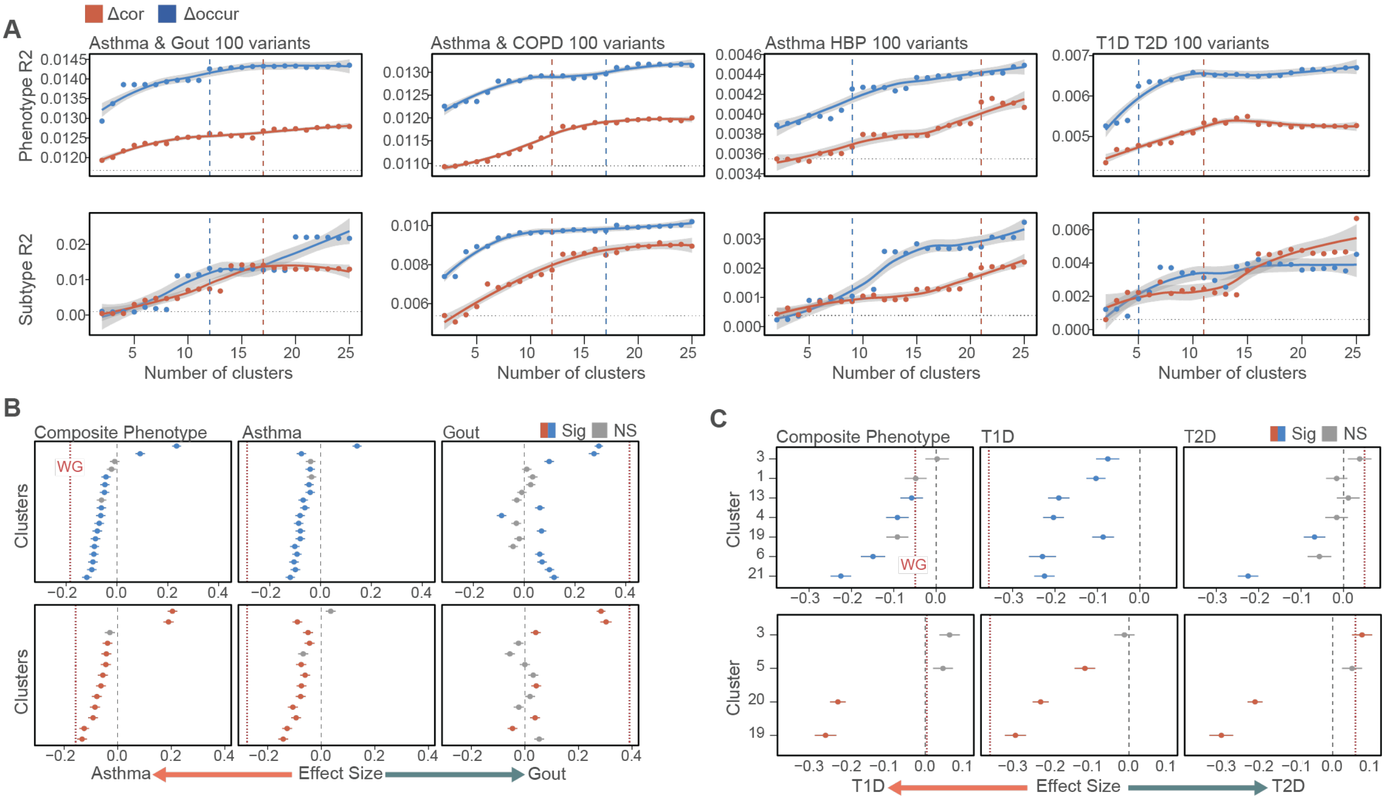


**Supplementary Figure 1. Application of variant cooccurrence to real composite phenotypes. (A)** Optimisation of the cluster number by the proportion of variance captured between composite cases and controls and alignment with the proportion of variance between composite cohort cases captured. Dashed lines represent the estimated plateau point. **(B-C)** Association of the best performing clustering iteration with each subtype when calculated using composite weights and subtype-specific weights for **(B)** asthma-gout and **(C)** T1D-T2D composite traits.
WG: whole genome.

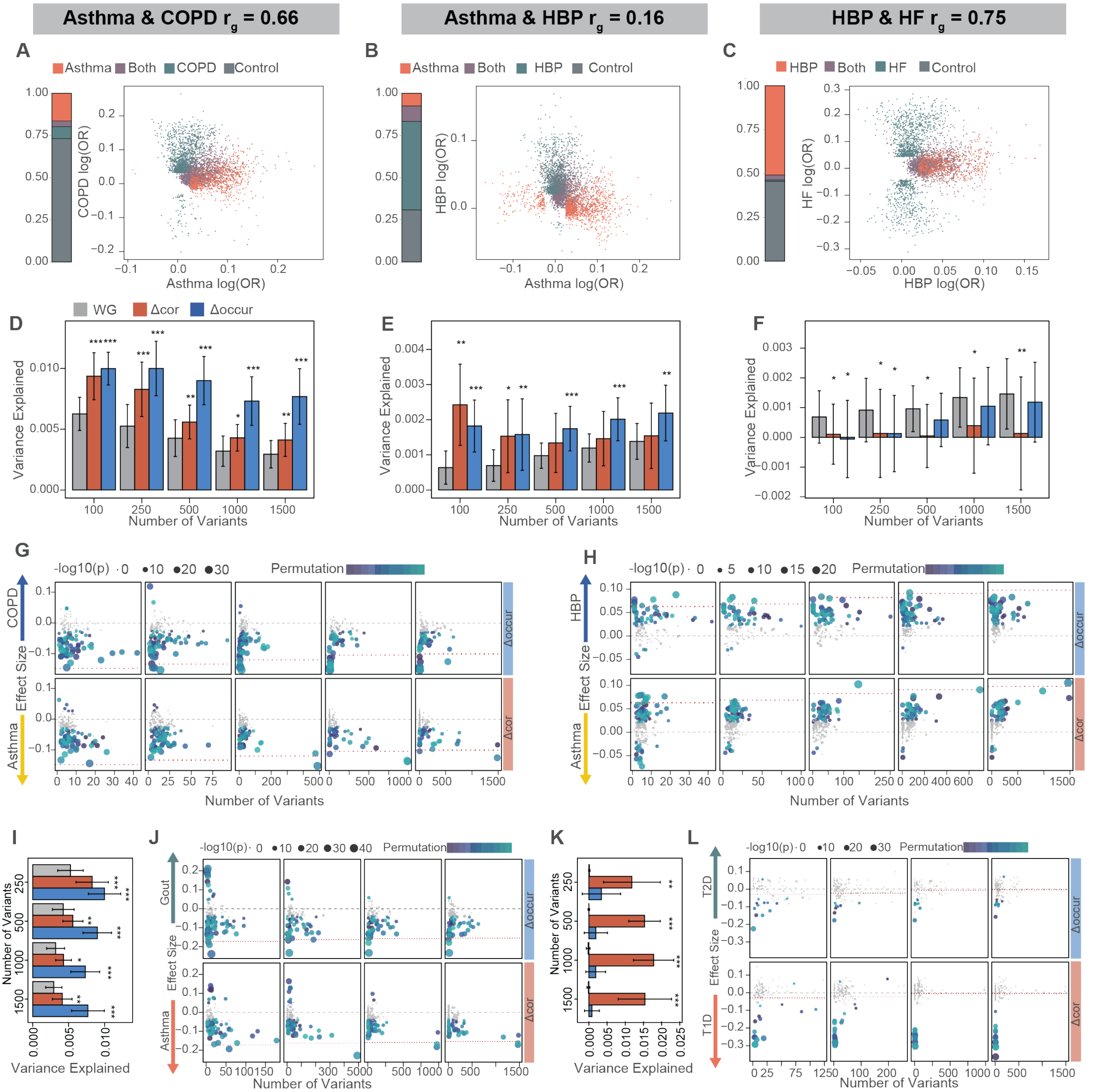


**Supplementary Figure 2. Validation of Δcor and Δoccur based approaches in additional composite phenotypes and across increasing numbers of variants. (A-C)** Cohort composition and GWAS associations of composite traits comprised of **(A)** asthma and COPD, **(B)** asthma and hypertension, **(C)** hypertension and heart failure. **(D-F)** Proportion of subtype variance (Nagelkerke's $R^2$) explained by jointly fitting all cluster-partitioned PRS compared to the unpartitioned PRS for each composite cohort across increasing numbers of variants. **(G-H)** Effect size of each cluster-PRS in separating disease subtypes across increasing numbers of variants summarised across 10 permutations. **(I-J) (I)** Proportion of variance captured by cluster-PRS and **(J)** effect size distribution of each cluster-PRS across increasing numbers of variants for asthma-gout. **(K-L)** as per (I-J) forT1D-T2D composite traits.
Error bars indicate standard deviation. p < 0.05 * p < 0.01 ** p < 0.001 ***.
HBP: high blood pressure, WG: Whole Genome.

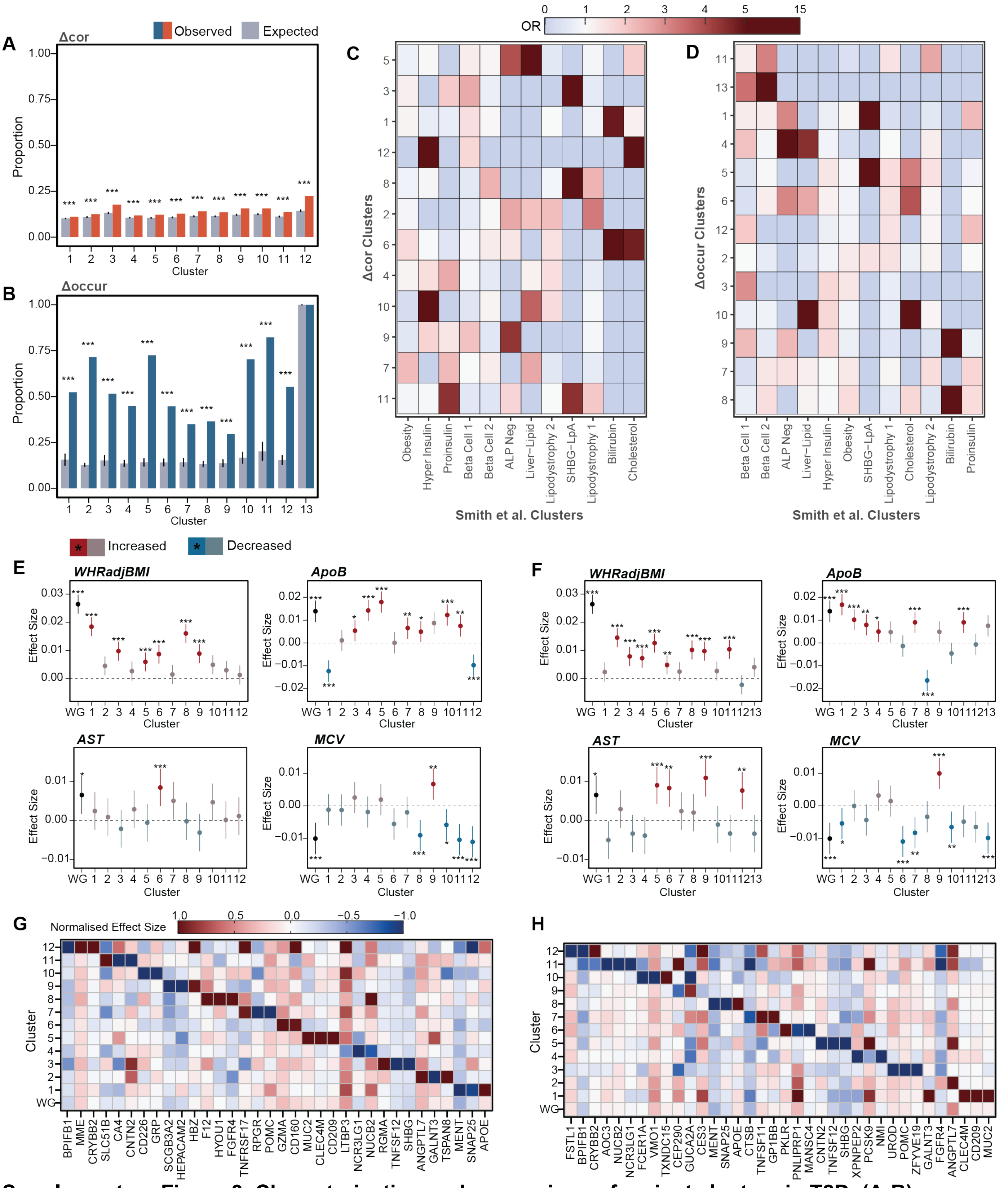


**Supplementary Figure 3. Characterisation and comparison of variant clusters in T2D. (A-B)** Conservation of clusters identified by variant **(A)** Δcor and **(B)** Δoccur with subsampling of the disease cohort compared to expectation by chance. **(C-D)** Enrichment for variant sharing between clusters identified by **(C)** Δcor and **(D)** Δoccur and clusters identified by Smith *et al.* (2024). **(E-F)** Association of cluster variants and cluster-partitioned PRS across key T2D risk factors and traits for **(E)** Δcor and **(F)** Δoccur derived clusters. **(G-H)** The top 3 associated plasma proteins per cluster for **(G)** Δcor and **(H)** Δoccur derived clusters.
Error bars indicate 95% confidence intervals. $p < 0.05$ * $p < 0.01$ ** $p < 0.001$ ***.
WG: Whole Genome.

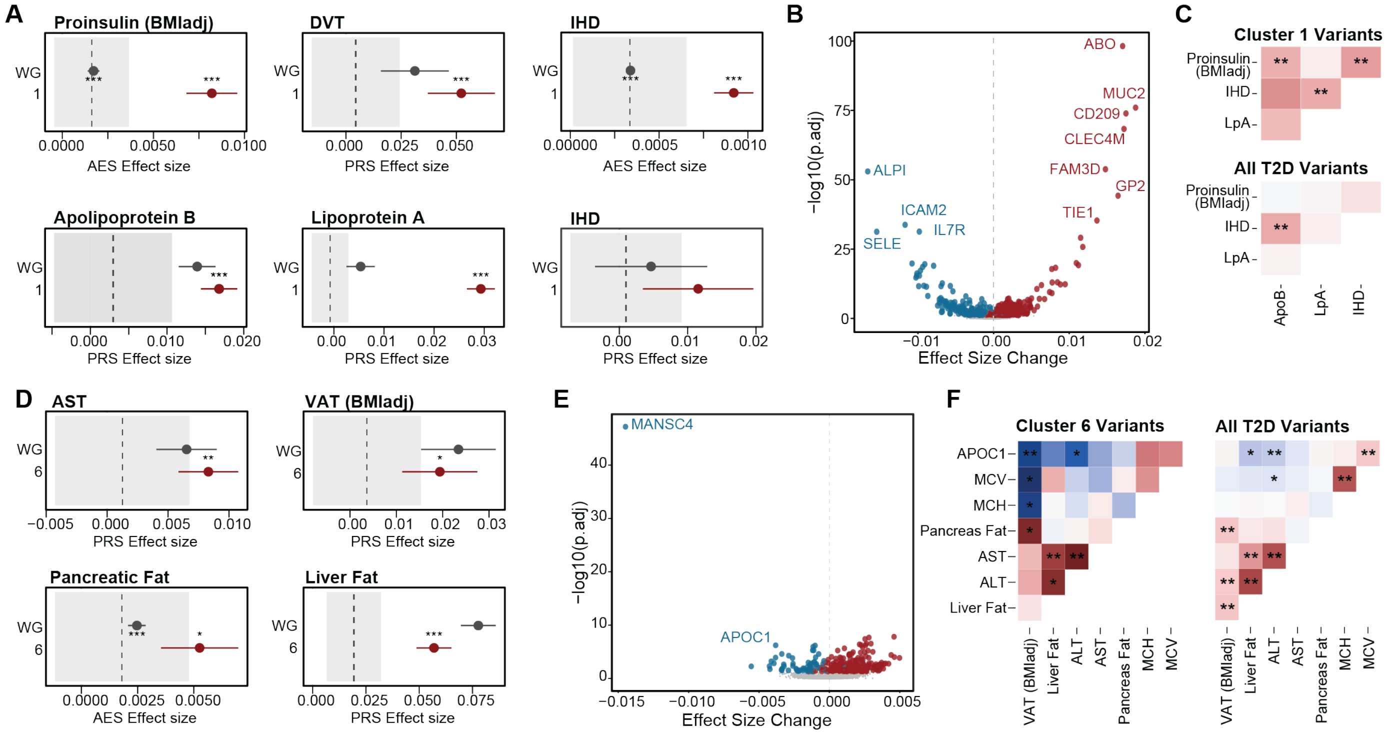


**Supplementary Figure 4. Cooccurrence derived clusters show shared genetic regulation of coherent mechanistic profiles. (A)** Associations for distinguishing traits for Δoccur cluster 1 (AES and PRS). The aggregated variant set (WG) is provided as reference. Dashed line with grey box shows the average ± standard deviation of all other cluster effect sizes. **(B)** Distinguishing plasma protein associations of cluster 1 compared to all variants. **(C)** Partitioned genetic correlation analysis between cluster 1 associated traits, partitioned by cluster 1 variants or all T2D variants. **(D-E)** Distinguishing **(D)** traits and **(E)** plasma proteins for Δoccur cluster 6. **(F)** Partitioned genetic correlation analysis between cluster-6 associated traits and biomarkers, partitioned by cluster 6 variants or all T2D variants.
WG: Whole genome. p < 0.05 * p < 0.01 ** p < 0.001 ***

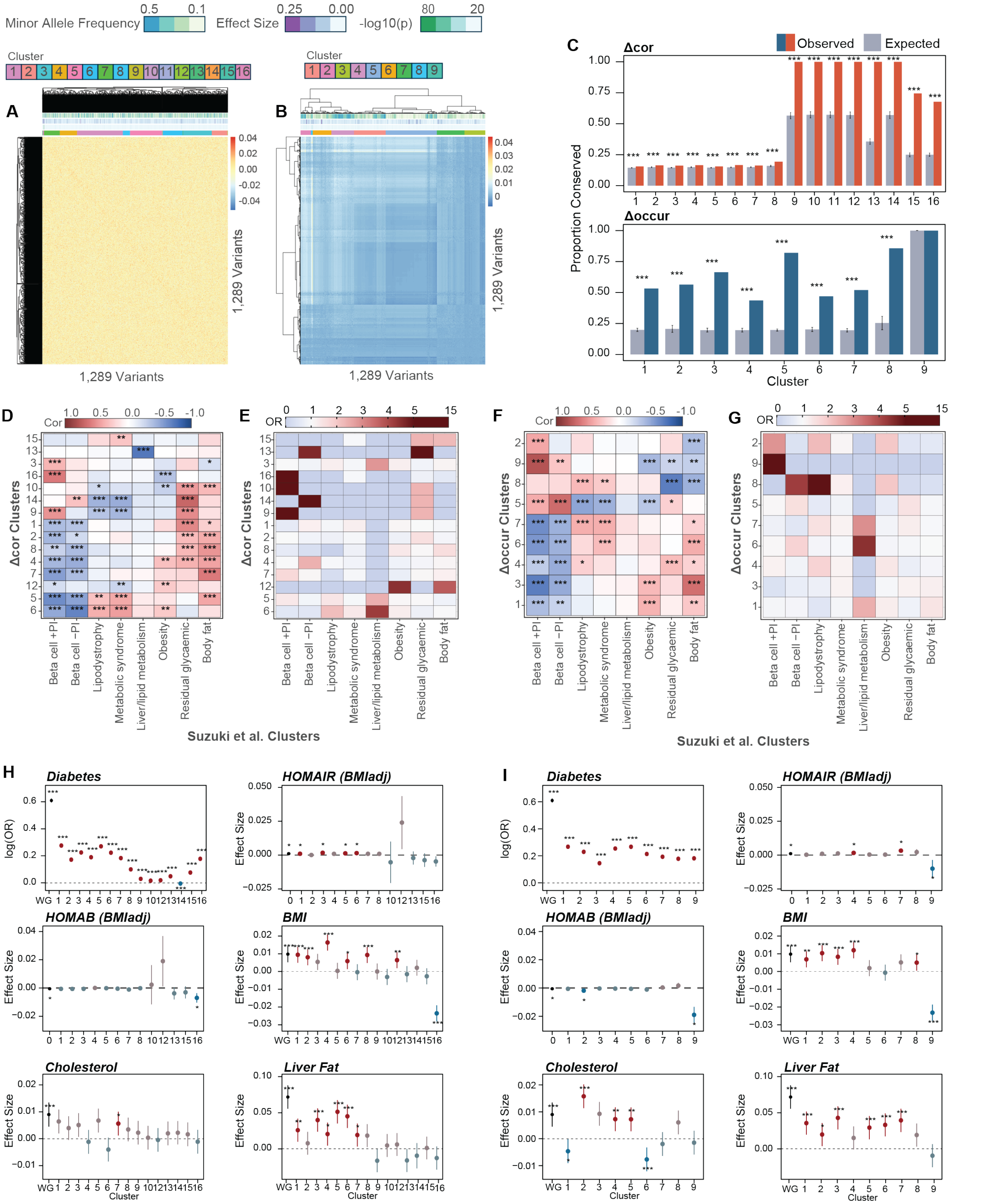


**Supplementary Figure 5. Characterisation of cooccurrence clusters derived from the Suzuki *et al.* variants for T2D. (A-B)** Variant-variant **(A)** Δcor and **(B)** Δoccur matrix of 1,289 T2D-associated variants divided annotated by effect size, minor allele frequency, p-value and cluster assignment. **(C)** Conservation of clusters identified by variant Δcor and Δoccur with subsampling of the disease cohort compared to expectation by chance. **(D-E)** Overlap of Δcor derived clusters by **(D)** AES across 123 phenotypes and **(E)** variant sharing. **(F-G)** Overlap of Δoccur derived clusters by **(D)** AES across 123 phenotypes and **(E)** variant sharing. **(H-I)** Association of cluster variants and cluster-partitioned PRS across key T2D risk factors and traits for **(H)** Δcor and **(I)** Δoccur derived clusters. Effect size shows change in standard deviation of trait per change in standard deviation of PRS.
Error bars indicate 95% confidence intervals. p < 0.05 * p < 0.01 ** p < 0.001 ***.

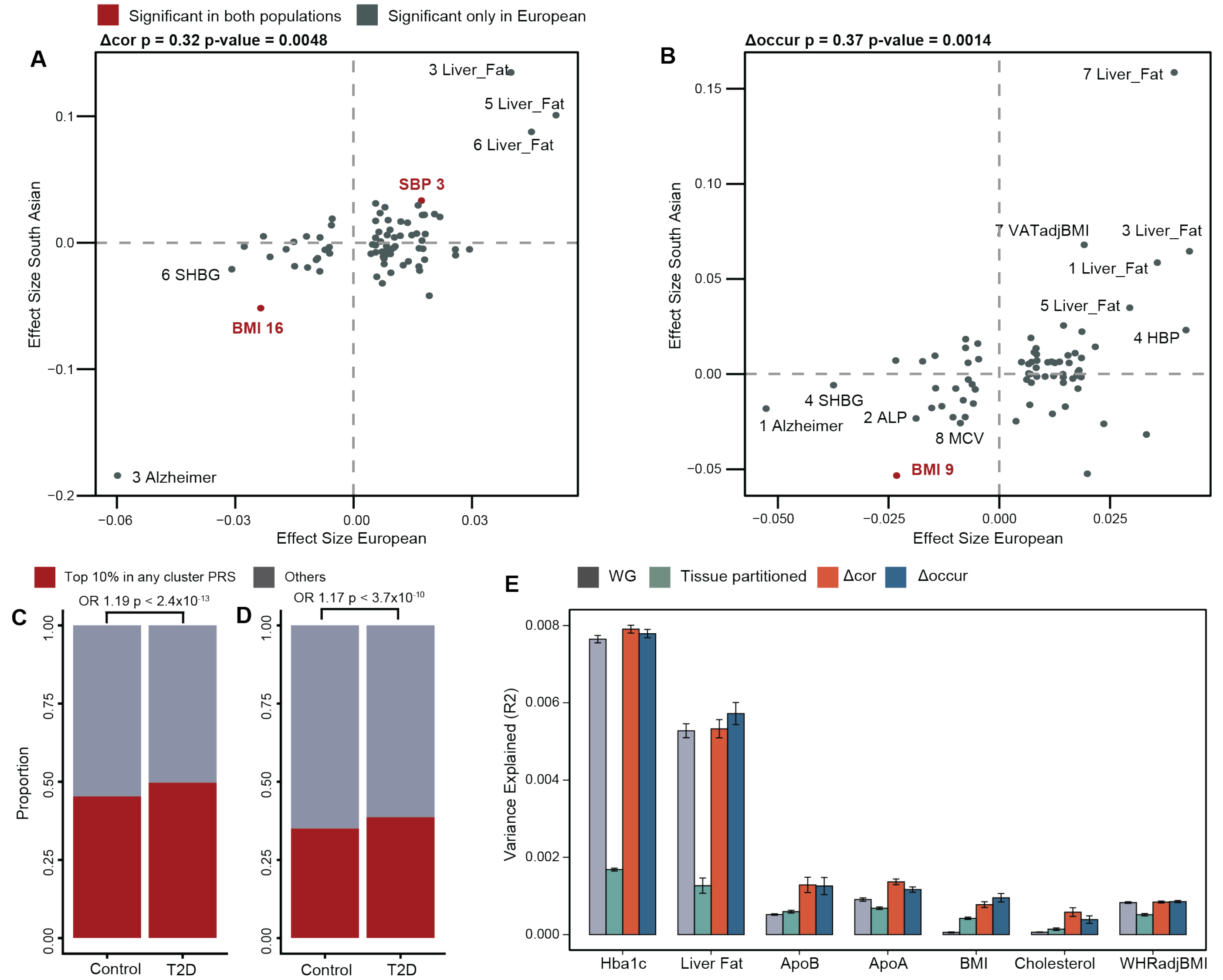


**Supplementary Figure 6. Validation and application of cooccurrence derived clusters in T2D using the Suzuki *et al* variant set.**

**(A-B).** Correlation of effect sizes for each cluster-PRS and phenotype pair between European and South Asian cohorts for **(A)** Δcor and **(B)** Δoccur derived clusters. Significant associations in the South Asian cohort ($p < 0.05$) are highlighted. Spearman's correlation coefficient reported. **(C-D)** Individuals with T2D but low overall genetic risk for T2D risk are detected in the top 10% of at least one cluster-partitioned PRS in both **(C)** Δcor and **(D)** Δoccur. Chi-squared enrichment test reported. **(E)** Proportion of phenotypic variance ($R^2$) explained by jointly fitting all cluster partitioned-PRS compared to the unpartitioned PRS and tissue-partitioned PRS across 10 permutations. Error bars show standard error. p < 0.05 * p < 0.01 ** p < 0.001 ***

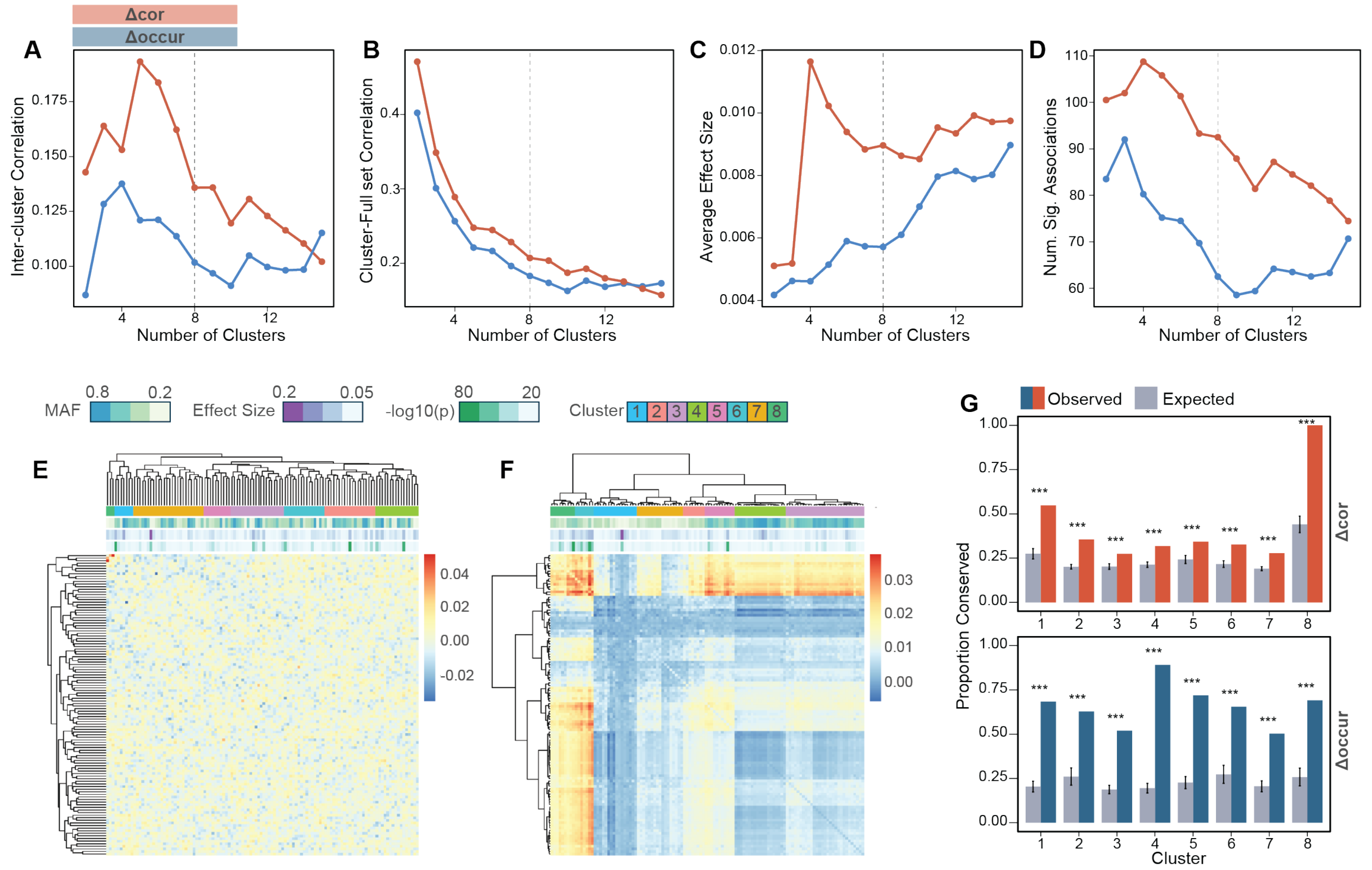


**Supplementary Figure 7. Optimization and benchmarking of variant sets by their functional distinctiveness. (A)** Average inter-cluster Jaccard correlation using the phenotype panel across increasing numbers of clusters. **(B)** Average Jaccard correlation of each cluster with the full variant set using the phenotype panel across increasing numbers of clusters. **(C)** Average absolute aggregated effect size in a panel of 93 phenotypes across increasing numbers of clusters. **(D)** Average number of significant associations per cluster in a panel of 93 phenotypes across increasing numbers of clusters. **(E-F)** Variant **(E)** Δcor and **(F)** Δoccur, annotated by cluster assignment, minor allele frequency, effect size and significance in asthma. **(G)** Conservation of clusters identified by variant Δcor and Δoccur with subsampling of the disease cohort compared to expectation by chance.

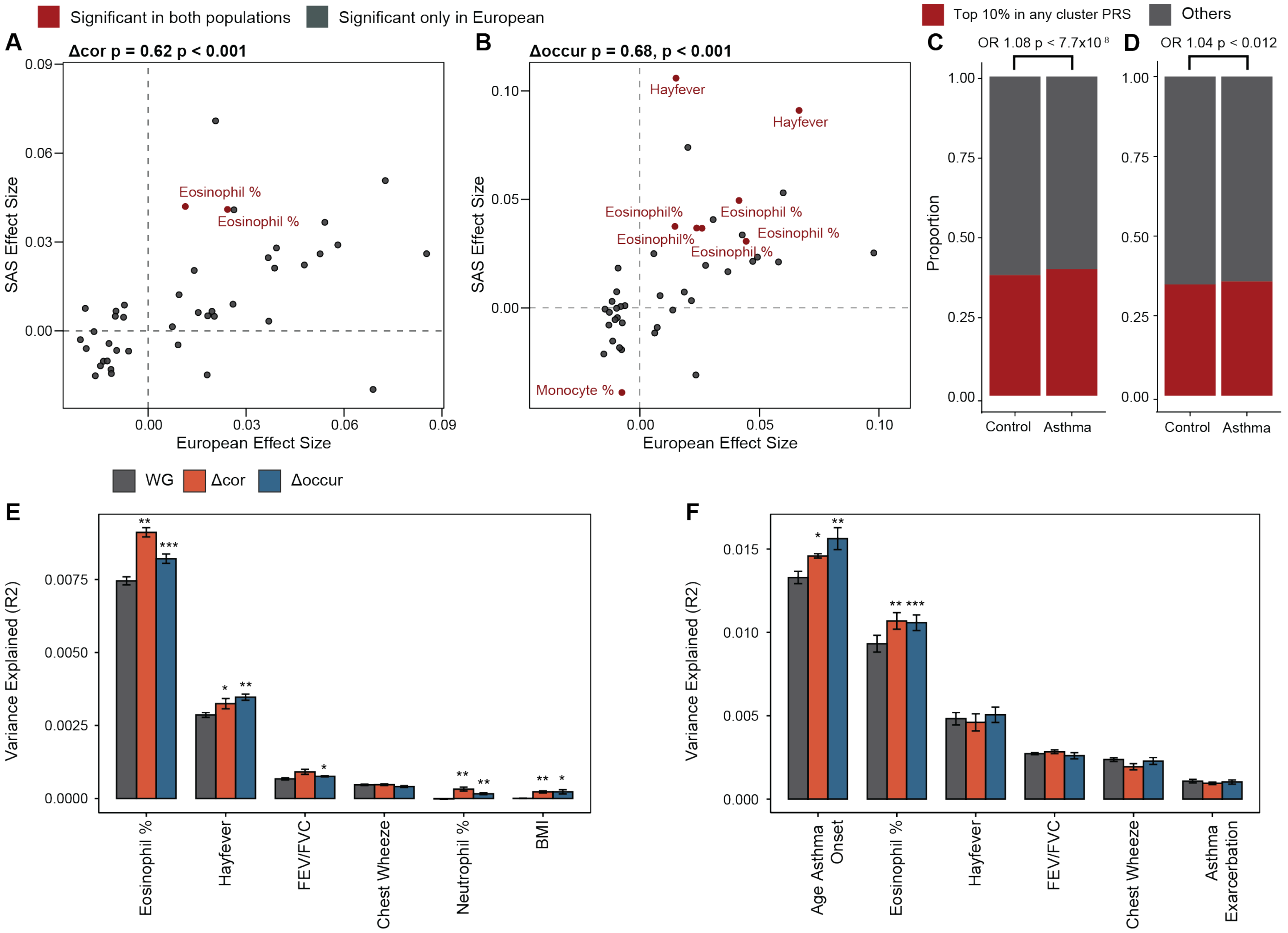


**Supplementary Figure 8. Validation and application of cooccurrence derived clusters in asthma. (A-B).** Correlation of effect sizes for each cluster-PRS and phenotype pair between European and South Asian cohorts for **(A)** Δcor and **(B)** Δoccur derived clusters. Significant associations in the South Asian cohort ($p < 0.05$) are highlighted. Spearman's correlation coefficient reported. **(C-D)** Individuals with asthma but low overall genetic risk for asthma risk is detected in the top 10% of at least one cluster-partitioned PRS in both **(C)** Δcor and **(D)** Δoccur. Chi-squared enrichment test reported. **(E-F)** Proportion of phenotypic variance ($R^2$) explained by jointly fitting all cluster partitioned-PRS compared to the unpartitioned PRS in the **(E)** full cohort and **(F)** asthma cohort across 10 permutations. Error bars show standard deviation.

p < 0.05 * p < 0.01 ** p < 0.001 ***

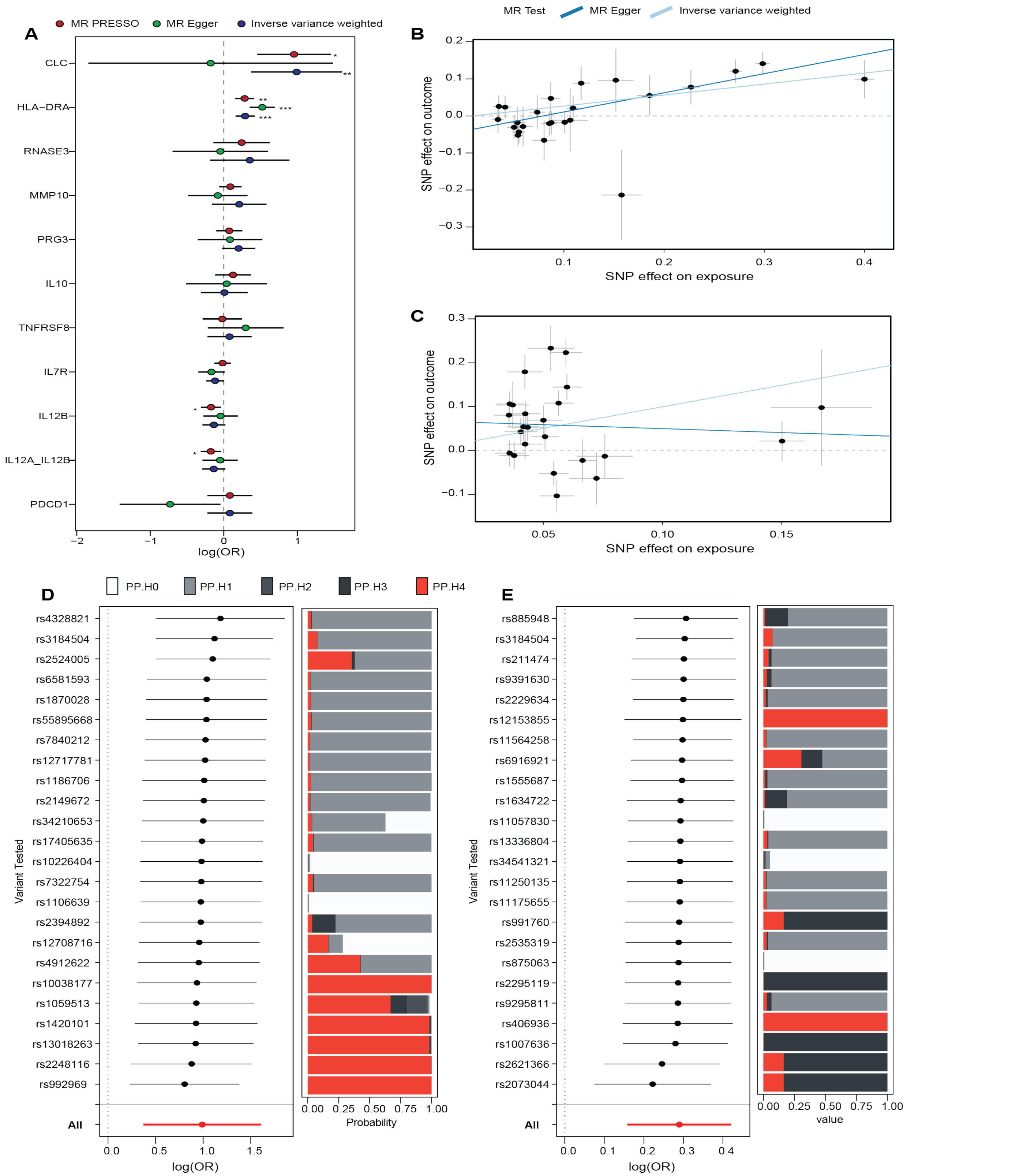


**Supplementary Figure 9. Mendelian Randomization analysis of plasma protein associated with asthma exacerbation. (A)** Summary of MR results for 10 candidate plasma proteins using MR PRESSO, MR Egger and inverse-variance weighted analysis. **(B-C)** Scatter plots showing correlation between effect size in the exposure **(B)** HLA-DRA or **(C)** CLC and outcome (asthma hospitalisation) with MR Egger and IVW estimates shown. **(D-E)** Leave one out and colocalization analysis evaluating the effect of each SNP on the overall MR estimate for **(B)** HLA-DRA or **(C)** CLC and the probability that the same SNP is causal for both protein abundance and asthma hospitalisation.
